# Learning accurate path integration in a ring attractor model of the head direction system

**DOI:** 10.1101/2021.03.12.435035

**Authors:** Pantelis Vafidis, David Owald, Tiziano D’Albis, Richard Kempter

## Abstract

Ring attractor models for angular path integration have recently received strong experimental support. To function as integrators, head-direction (HD) circuits require precisely tuned connectivity, but it is currently unknown how such tuning could be achieved. Here, we propose a network model in which a local, biologically plausible learning rule adjusts synaptic efficacies during development, guided by supervisory allothetic cues. Applied to the *Drosophila* HD system, the model learns to path-integrate accurately and develops a connectivity strikingly similar to the one reported in experiments. The mature network is a quasi-continuous attractor and reproduces key experiments in which optogenetic stimulation controls the internal representation of heading, and where the network remaps to integrate with different gains. Our model predicts that path integration requires supervised learning during a developmental phase. The model setting is general and also applies to architectures that lack the physical topography of a ring, like the mammalian HD system.

## Introduction

Spatial navigation is crucial for the survival of animals in the wild and has been studied in many model organisms (Tolman, 1948; O’Keefe et al., 1978; Gallistel, 1993; Eichenbaum, 2017). To orient themselves in an environment, animals rely on external sensory cues (e.g. visual, tactile, or auditory), but such allothetic cues are often ambiguous or absent. In these cases, animals have been found to update internal representations of their current location based on idiothetic cues, a process that is termed path integration (PI, Darwin, 1873; Mittelstaedt and Mittelstaedt, 1980; McNaughton et al., 1996; Etienne et al., 1996; Burak and Fiete, 2009). The head direction (HD) system is a simple example of a circuit that can support PI, and head direction cells in rodents and flies provide an internal representation of orientation that can persist in darkness (Ranck, 1984; Mizumori and Williams, 1993; Seelig and Jayaraman, 2015).

In rodents, the internal representation of heading takes the form of a localized “bump” of activity in the high-dimensional neural manifold of HD cells (Chaudhuri et al., 2019). It has been proposed that such a localized activity bump could be sustained by a ring attractor network with local excitatory connections (Skaggs et al., 1995; Redish et al., 1996; Hahnloser, 2003; Samsonovich and McNaughton, 1997; Song and Wang, 2005; Stringer et al., 2002; Xie et al., 2002), resembling reverberation mechanisms proposed for working memory (Wang, 2001). Ring attractor networks used to model HD cells fall in the theoretical framework of continuous attractor networks (CANs, Amari, 1977; Ben-Yishai et al., 1995; Seung, 1996). In this setting, HD cells can update the heading representation even in darkness by smoothly moving the bump around the ring obeying idiothetic angular-velocity cues.

Interestingly, a physical ring-like attractor network of HD cells was demonstrated in the *Drosophila* central complex (CX, Seelig and Jayaraman, 2015; Green et al., 2017, 2019; Franconville et al., 2018; Kim et al., 2019; Fisher et al., 2019; Turner-Evans et al., 2020). Notably, in *Drosophila* (from here on simply referred to as “fly”), HD cells (named E-PG neurons, also referred to as “compass” neurons) are physically arranged in a ring, and an activity bump is readily observable from a small number of cells (Seelig and Jayaraman, 2015). Moreover, as predicted by some computational models (Skaggs et al., 1995; Samsonovich and McNaughton, 1997; Stringer et al., 2002; Song and Wang, 2005), the fly HD system also includes cells (named P-EN1 neurons) that are conjunctively tuned to head direction and head angular velocity. We refer to these neurons as head rotation (HR) cells because of their putative role in shifting the HD bump across the network according to the head’s angular velocity (Turner-Evans et al., 2017, 2020). One of the challenges in using such a model for PI is to sustain a bump of activity and move it with the right speed and direction around the ring.

Ring attractor models that act as path integrators require that synaptic connections are precisely tuned (Hahnloser, 2003). Therefore, if the circuit was completely hardwired, the amount of information that an organism would need to genetically encode connection strenghts would be exceedingly high. Additionally, it would be unclear how these networks could cope with variable sensory experiences. In fact, remarkable experimental studies have shown that when animals are placed in a virtual reality environment where visual and self-motion information can be manipulated independently (Stowers et al., 2017), PI capabilities adapt accordingly (Jayakumar et al., 2019). These findings suggest that PI networks are able to self-organize and to constantly recalibrate.

Here, we propose that a simple local learning rule could support the emergence of a PI circuit during development and its re-calibration once the circuit has formed. Specifically, we suggest that accurate PI is achieved by associating allothetic and idiothetic inputs at the cellular level. When available, the allothetic sensory input (here chosen to be visual) acts as a “teacher” signal in a setup that resembles supervised learning. The learning rule then exploits the relation between the allothetic heading of the animal (given by the visual input) and the idiothetic self-motion cues (which are always available), to learn how to integrate the latter.

The learning rule is inspired by previous experimental and computational work on mammalian cortical pyramidal neurons, which are believed to associate inputs to different compartments through an in-built cellular mechanism (Larkum, 2013; Urbanczik and Senn, 2014; Brea et al., 2016). In fact, it was recently shown that in layer 5 pyramidal cells internal and external information about the world arrive at distinct anatomical locations, and active dendritic gating controls learning between the two (Doron et al., 2020). In a similar fashion, we propose that learning PI in the HD system occurs by associating inputs at opposite poles of compartmentalized HD neurons, which we call “associative neurons” (Urbanczik and Senn, 2014; Brea et al., 2016).

In summary, here we show for the first time how a biologically plausible synaptic plasticity rule enables to learn and maintain the complex circuitry required for PI. We apply our framework to the fly HD system because it is well characterized; yet our model setting is general and can be used to learn PI in other animal models once more details about the HD circuit there are known (Abbott et al., 2020). We find that the learned network is a ring attractor with a connectivity that is strikingly similar to the one found in the fly CX (Turner-Evans et al., 2020) and that it can accurately path-integrate in darkness for the entire range of angular velocities that the fly displays. Crucially, the learned network accounts for several key findings in the experimental literature, and it generates testable predictions.

## Results

To illustrate basic principles of how PI could be achieved, we study a computational model of the HD system and show that synaptic plasticity could shape its circuitry through visual experience. In particular, we simulate the development of a network that, after learning, provides a stable internal representation of head direction and uses only angular-velocity inputs to update the representation in darkness. The internal representation of heading (after learning) takes the form of a localized bump of activity in the ring of HD cells. All neurons in our model are rate-based, i.e., spiking activity is not modeled explicitly.

## Model setup

The gross model architecture closely resembles the one found in the fly CX (Fig. 1A). It comprises HD cells organized in a ring, and HR cells organized in two wings. One wing is responsible for leftward and the other for rightward movement of the internal heading representation. HD cells receive visual input from the so-called “ring” neurons; this input takes the form of a disinhibitory bump centered at the current HD (Fig. 1B and eq. (4), Omoto et al., 2017; Fisher et al., 2019). The location of this visual bump in the network is controlled by the current head direction. We simulate head movements by sampling head-turning velocities from an Ornstein–Uhlenbeck process (Methods), and we provide the corresponding velocity input to the HR cells (eq. (9), Fig. 1C). HR cells provide direct input to HD cells, and HR cells also receive input from HD cells (Fig. 1A). Both HR and HD cells receive global inhibition, which is in line with a putative “local” model of HD network organization (Kim et al., 2017). The connections from HR to HD cells (*W*^*HR*^) and the recurrent connections among HD cells (*W*^*rec*^) are assumed to be plastic. The goal of learning is to tune these plastic connections so that the network can achieve PI in the absence of visual input.

**Figure 1.**
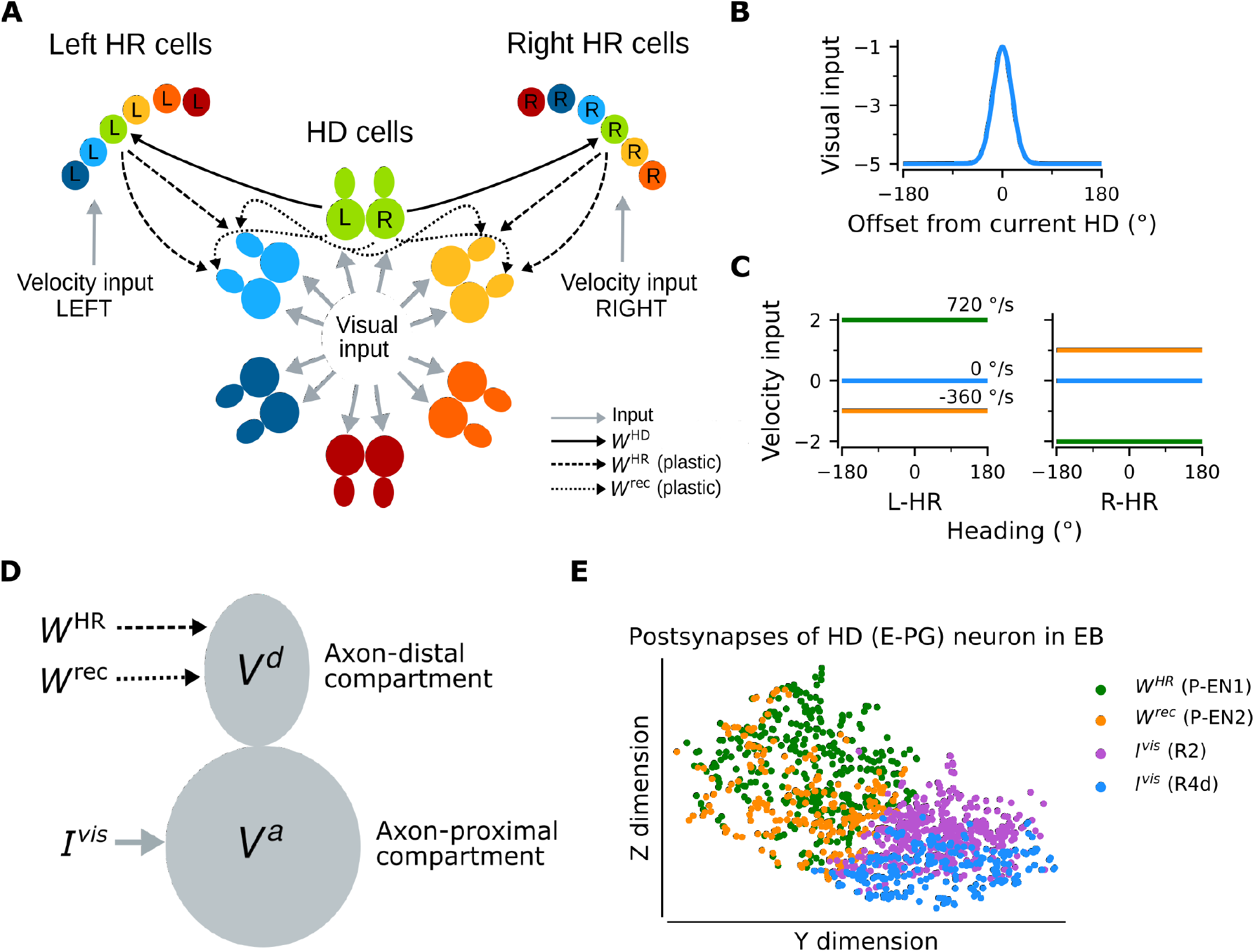
Network architecture. (A) The ring of HD cells projects to two wings of HR cells, a leftward (Left HR cells, abbreviated as L-HR) and a rightward (Right HR cells, or R-HR), so that each wing receives selective connections only from a specific HD cell (L: left, R: right) for every head direction. For illustration purposes, the network is scaled-down by a factor of 5 compared to the original cell numbers *N*^*HR*^ = *N*^*HD*^ = 60. The schema shows the outgoing connections (*W*^*HD*^ and *W*^*rec*^) only from the green HD neurons and the incoming connections (*W*^*HR*^ and *W*^*rec*^) only to the light blue and yellow HD neurons. Furthermore, the visual input to HD cells and the velocity inputs to HR cells are indicated. Visual input to the ring of HD cells as a function of radial distance from the current head direction. Angular-velocity input to the wings of HR cells for three angular velocities. (D) The associative neuron: *V*^*a*^ and *V*^*d*^ denote the voltage in the axon-proximal (i.e. closer to the axon initial segment) and axon-distal (i.e. further away from the axon initial segment) compartment, respectively. Arrows indicate the inputs to the compartments, as in (A), and *I*^*vis*^ is the visual input current. (E) Postsynaptic locations in the ellipsoid body (EB) for an example HD (E-PG) neuron; for details, see Methods. The neuron receives recurrent and HR input (green and orange dots, corresponding to inputs from P-EN1 and P-EN2 cells, respectively) and visual input (purple and blue dots, corresponding to inputs from visually responsive R2 and R4d cells, respectively) in distinct spatial locations.

The unit that controls plasticity in our network is an “associative neuron”. It is inspired by pyramidal neurons of the mammalian cortex whose dendrites act, via backpropagating action potentials, as coincidence detectors for signals arriving from different layers of the cortex and targeting different compartments of the neuron (Larkum et al., 1999). Paired with synaptic plasticity, coincidence detection can lead to long-lasting associations between these signals (Larkum, 2013). To map the morphology of a cortical pyramidal cell to the one of a HD cell in the fly, we first point out that all relevant inputs arrive at the dendrites of HD cells within the ellipsoid body (EB) of the fly (Xu et al., 2020); moreover, the soma itself is externalized in the fly brain, and it is unlikely to contribute considerably to computations (Gouwens and Wilson, 2009; Tuthill, 2009). We thus link the dendrites of the pyramidal associative neuron to the axon-distal dendritic compartment of the associative HD neuron in the fly, and we link the soma of the pyramidal associative neuron to the axon-proximal dendritic compartment of the associative HD neuron in the fly. Furthermore, we assume that the axon-proximal compartment is electrotonically closer to the axon initial segment, and therefore, similarly to the somatic compartment in pyramidal neurons, inputs there can more readily initiate action potentials. We also assume that associative HD cells receive visual input (*Ivis*) in the axon-proximal compartment, and both recurrent input (*W*^*rec*^) and HR input (*W*^*HR*^) in the axon-distal compartment; accordingly, we model HD neurons as two-compartment units (Fig. 1D). The associative neuron can learn the synaptic weights of the incoming connections in the axon-distal compartment, therefore, as mentioned, we let *W*^*rec*^ and *W*^*HR*^ be plastic.

We find that the assumption of spatial segregation of postsynapses of HD cells is consistent with our analysis of recently-released EM data from the fly (Xu et al., 2020). For an example HD (E-PG) neuron, Figure 1E depicts that head rotation and recurrent inputs (mediated by P-EN1 and P-EN2 cells, respectively (Turner-Evans et al., 2020)) contact the E-PG cell in locations within the EB that are distinct compared to those of visually responsive neurons R2 and R4d (Omoto et al., 2017; Fisher et al., 2019), as hypothesized. The same pattern was observed for a total of 16 E-PG neurons (one for each “wedge” of the EB) that we analyzed.

The connections from HD to HR cells (*W*^*HD*^) are assumed to be fixed, and HR cells are modeled as single-compartment units. Projections are organized such that each wing neuron receives input from only one specific HD neuron for every HD (Fig. 1A). This simple initial wiring makes HR cells conjunctively tuned to HR and HD, and we assume that it has already been formed, for example, during pre-natal circuit assembly. In addition, the connections carrying the visual and angular velocity inputs are also assumed to be fixed. Although plasticity in the visual inputs has been shown to exist (Fisher et al., 2019; Kim et al., 2019), here we focus on how the path-integrating circuit itself originally self-organizes. Therefore, to simplify the setting and without loss of generality, we assume a fixed anchoring to environmental cues as the animal moves in the same environment (for details, see Discussion).

The visual input acts as a supervisory signal during learning (D’Albis and Kempter, 2020), which is used to change weights of synapses onto the axon-distal compartment of HD cells. We utilize the learning rule proposed by Urbanczik and Senn (2014) (for details, see Methods), which tunes the incoming synaptic connections in the axon-distal compartment in order to minimize the discrepancy between the firing rate of the neuron (which is primarily a function of the visual input) and the prediction of the firing rate by the axon-distal compartment in the absence of visual input, which depends on head rotation velocity. From now on, we refer to this discrepancy as “learning error”, or simply “error” (eq. (17)) (in units of firing rate). Importantly, this learning rule is biologically plausible because the firing rate of an associative neuron is locally available at every synapse in the axon-distal compartment due to the assumed back-propagation of axonal activity to the dendrites. The other two signals that enter the learning rule are the voltage of the axon-distal compartment and the postsynaptic potential, which are also available locally at the synapse; for details, see Methods.

### Mature network can path-integrate in darkness

Figure 2A shows an example of the performance of a trained network, for the light condition (i.e., when visual input is available; yellow overbars) and for PI in darkness (purple overbars); the performance is quantified by the PI error (in units of degrees) over time. PI error refers to the accumulated difference between the internal representation of heading and the true heading, and it is different from the learning error introduced previously.

**Figure 2.**
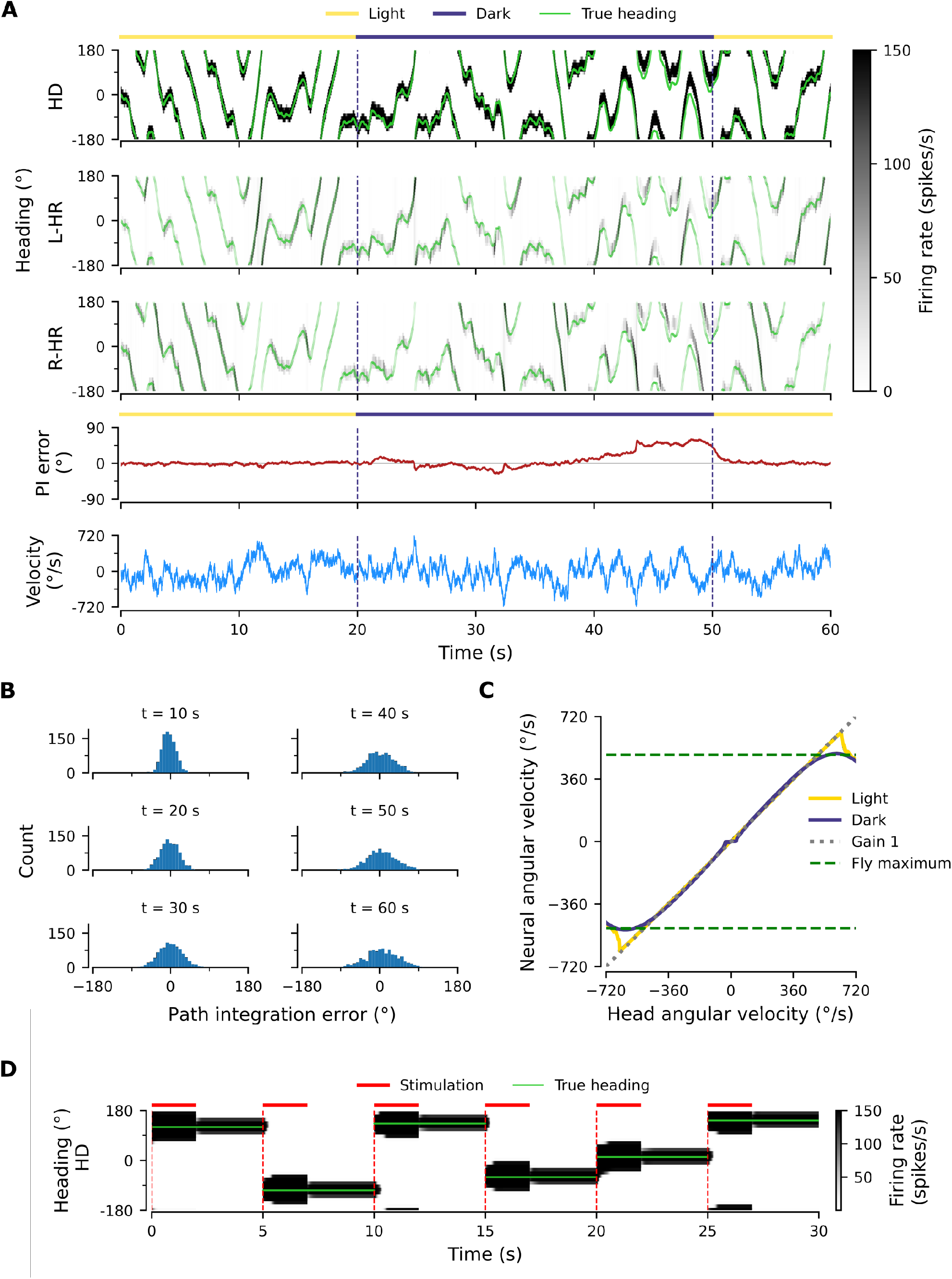
Path integration (PI) performance of the network. (A) Example activity profiles of HD, L-HR, and R-HR neurons (firing rates gray-scale coded). Activities are visually guided (yellow overbars) or are the result of PI in the absence of visual input (purple overbar). The ability of the circuit to follow the true heading is slightly degraded during PI in darkness. The PI error, i.e., the difference between the PVA and the true heading of the animal as well as the instantaneous head angular velocity are plotted separately. (B) Temporal evolution of the distribution of PI errors in darkness, for 1000 simulations. The distribution gets wider with time, akin to a diffusion process. (C) Relation between head angular velocity and neural angular velocity, i.e., the speed with which the bump moves in the network. There is almost perfect (gain 1) PI in darkness for head angular velocities within the range of maximum angular velocities that are displayed by the fly (dashed green horizontal lines; see Methods). (D) Example of consecutive stimulations in randomly permeated HD locations, simulating optogenetic stimulation experiments in Kim et al. (2017). Red overbars indicate when the network is stimulated with stronger than normal visual-like input, at the location indicated by the animal’s true heading (light green line), while red dashed vertical lines indicate the onset of the stimulation. The network is then left in the dark. Our simulations show that the bump remains at the stimulated positions, which suggests that the network well approximates a line attractor.

A unique bump of activity is clearly present at all times in the HD network (Fig. 2A, top), in both light and darkness conditions, and this bump moves smoothly across the network for a variable angular velocity (Fig. 2A, bottom). The position of the bump is defined as the population vector average (PVA) of the neural activity in the HD network. The HD bump also leads to the emergence of bumps in the HR network, separately for L-HR and R-HR cells (Fig. 2A, second and third panel from top). In light conditions (0–20 s in Fig. 2A), the PVA closely tracks the head direction of the animal in HD, L-HR, and R-HR cells alike, which is expected because the visual input guides the network activity. Importantly, however, in darkness (20–50 s in Fig. 2A), the self-motion input alone is enough to track the animal’s heading, leading to a small PI error between the internal representation of heading and the ground truth. This error is corrected after the visual input reappears (at 50 s in Fig. 2a). Such PI errors in darkness are qualitatively consistent with data reported in the experimental literature (Seelig and Jayaraman, 2015). The correction of the PI error also reproduces in silico the experimental finding that the visual input (whenever available) exerts stronger control on the bump location than the self-motion input (Seelig and Jayaraman, 2015), which suggests that even the mature network does not rely on PI when visual cues are available.

To quantify the accuracy of PI in our model, we draw 1000 trials, each 60 s long, for constant synaptic weights and in the absence of visual input. We also limit the angular velocities in these trials to retain only velocities that flies realistically display (see dashed green lines in Fig. 2C and Methods). We then plot the distribution of PI errors every 10 s (Fig. 2B). We find that average absolute PI errors (widths of distributions) increase with time in darkness, but most of the PI errors at 60 s are within 60 degrees of the true heading. This vastly exceeds the PI performance of flies (Seelig and Jayaraman, 2015). However, it should be noted that the model here corresponds to an ideal scenario which serves as a proof of principle. We will later incorporate irregularities owning to biological factors (asymmetry in the weights, biological noise) that bring the network’s performance closer to the fly’s behavior.

To further assess the network’s ability to integrate different angular velocities, we simulate the system both with and without visual input for 5 s during which the angular velocity is constant. We then compute the average movement velocity of the bump across the network, i.e., the neural velocity, and compare it to the real velocity provided as input. Figure 2C shows that the network achieves a PI gain (defined as the ratio between neural and real velocity) close to 1 both with and without supervisory visual input, meaning that the neural velocity matches very well the angular velocity of the animal, for all angular velocities that are observed in experiments (*v <* 500 degrees/s for walking and flying) (Geurten et al., 2014; Stowers et al., 2017). Although expected in light conditions, the fact that gain 1 is achieved in darkness shows that the network predicts the missing visual input from the velocity input, i.e., the network path integrates accurately. Note that PI is impaired in our model for very small angular velocities (Fig. 2C, flat purple line for *v <* 30 degrees/s), similarly to previous hand-tuned theoretical models (Turner-Evans et al., 2017). This is a direct consequence of the fact that maintaining a stable activity bump and moving it across the network at very small angular velocities are competing goals. Crucially, it has been reported that such an impairment of PI for small angular velocities exists in flies (Seelig and Jayaraman, 2015). Therefore our network reproduces this feature of the fly HD system as an emergent property from learning, and not as a feature built-in by hand.

### The network is a quasi-continuous attractor

A continuous attractor network (CAN) should be able to maintain a localised bump of activity in virtually a continuum of locations around the ring of HD cells. To prove that the learned network approximates this property, we seek to reproduce in silico experimental findings in Kim et al. (2017). There it was shown that local optogenetic stimulation of HD cells in the ring can cause the activity bump to jump to a new position and persist in that location — supported by internal dynamics alone.

To reproduce the experiments by Kim et al. (2017), we simulate optogenetic stimulation of HD cells in our network as visual input of increased strength and extent (for details, see Methods). We find that the strength and extent of the stimulation needs to be increased relative to that of the visual input; only in this case, a bump at some other location in the network can be suppressed, and a new bump emerges at the stimulated location. The stimuli are assumed to appear instantaneously at random locations, but we restrict our set of stimulation locations to the discrete angles represented by the finite number of HD neurons. Furthermore, the velocity input is set to zero for the entire simulation, signaling lack of head movement.

Figure 2D shows network activity in response to several stimuli, when the stimulation location changes abruptly every 5 s. During stimulation (2 s long, red overbars), the bump is larger than normal due to the use of a stronger than usual visual-like input to mimic optogenetic stimulation. The way in which the network responds to a stimulation depends on how far away from the “current” location it is stimulated: for shorter distances, the bump activity shifts to the new location, as evidenced by the transient dynamics at the edges of the bump resembling an exponential decay from an initial to a new location (e.g. see Fig. 2D at 20 s). However, for longer distances the bump first emerges in the new location and subsequently disappears at the initial location, a mechanism akin to a “jump” (Fig. 2D, all other transitions). Similar effects have been observed in the experimental literature (Seelig and Jayaraman, 2015; Kim et al., 2017). The way the network responds to stimulation indicates that it operates in a CAN manner, and not as a winner-takes-all (WTA) network where changes in bump location would always be instantaneous (Carpenter and Grossberg, 1987; Itti et al., 1998; Wang, 2002).

Following a 2-s stimulation, the network activity has converged to the new cued location. After the stimulation has been turned off, the bump remains at the new location (within the angular resolution Δ*ϕ* of the network), supported by internal network dynamics alone (Fig. 2D). We confirmed in additional simulations that the bump does not drift away from the stimulated location for extended periods of time (3-minute duration tested, only 3 s shown), and for all discrete locations in the HD network (only six locations shown). Therefore, we conclude that the HD network is a quasi-continuous attractor that can reliably sustain a heading representation over time in all HD locations. In reality, it is expected for the bump to drift away due to asymmetries in the connectivity of the biological circuit and biological noise (Burak and Fiete, 2012). In flies, for instance, the bump can stay put only for several seconds (Kim et al., 2017).

### Learning results in synaptic connectivity that matches the one in the ly

To gain more insight into how the network achieves PI and attains CAN properties, we show how the synaptic weights of the network are tuned during a developmental period (Fig. 3). Figures 3A,B show the learned recurrent synaptic weights among the HD cells, *W*^*rec*^, and the learned synaptic weights from HR to HD cells, *W*^*HR*^, respectively. Circular symmetry is apparent in both matrices, a crucial property for any ring attractor. Therefore we also plot the profiles of the learned weights as a function of receptive field difference in Fig. 3C.

**Figure 3.**
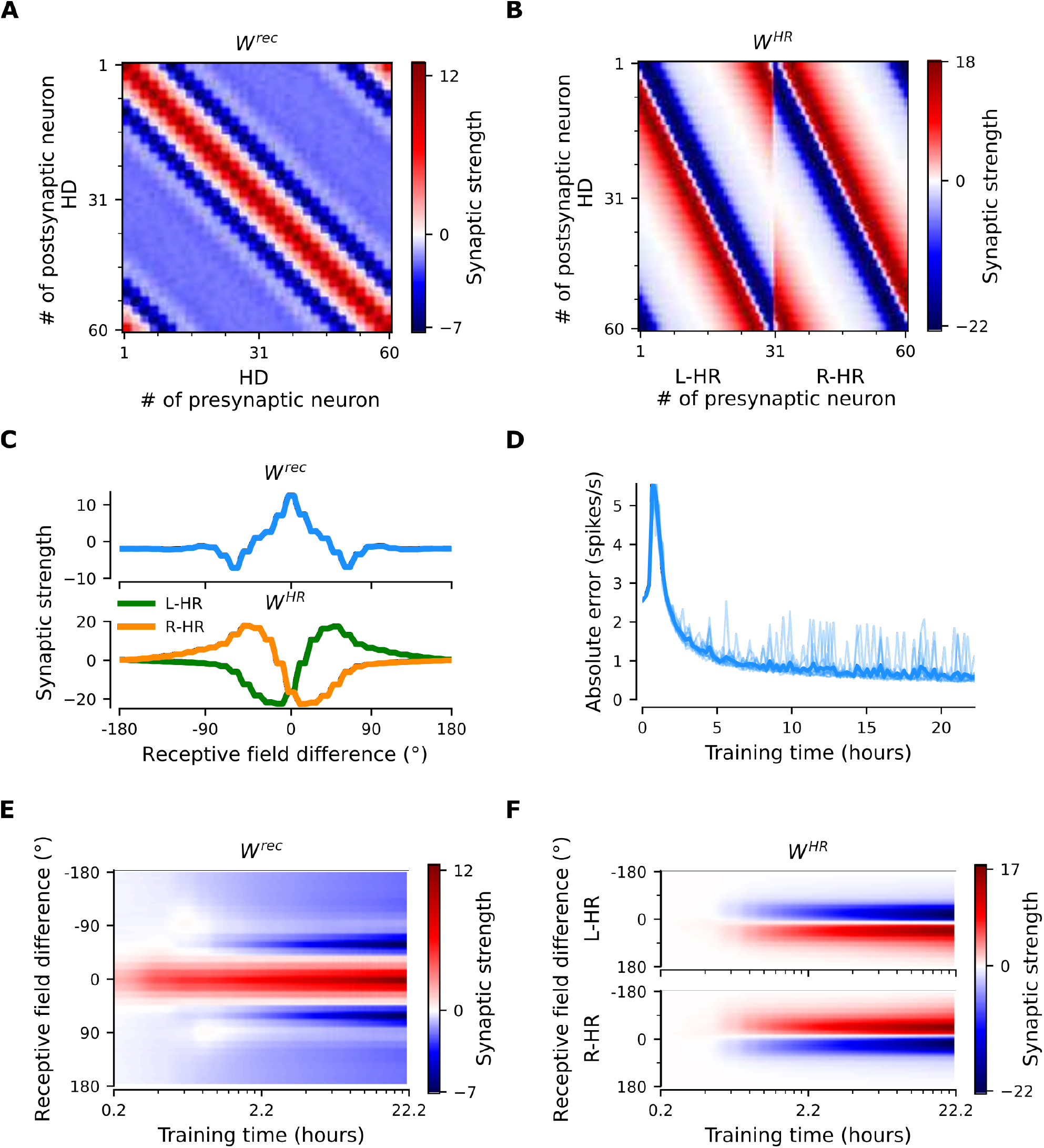
The network connectivity during and after learning. (A), (B) The learned weight matrices (color coded) of recurrent connections in the HD ring, *W*^*rec*^, and of HR-to-HD connections, *W*^*HR*^, respectively. Note the circular symmetry in both matrices. (C) Profiles of (A) and (B), averaged across presynaptic neurons. (D) Absolute learning error in the network (eq. (18)) for 12 simulations (transparent lines) and average across simulations (opaque line). At time *t* = 0, we initialize all the plastic weights at random and train the network for 8 × 10^4^ s (∼22 hours). The mean learning error increases in the beginning while a bump in *W*^*rec*^ is emerging, which is necessary to generate a pronounced bump in the network activity. For weak activity bumps, absolute errors are small because the overall network activity is low. After ∼1 hour of training, the mean learning error decreases with increasing training time and converges to a small value. (E), (F) Time courses of development of the profiles of *W*^*rec*^ and *W*^*HR*^, respectively. Note the logarithmic time scale.

First, we discuss the properties of the learned weights. Local excitatory connections have developed along the main diagonal of *W*^*rec*^, similar to what is observed in the CX (Turner-Evans et al., 2020). This local excitation can be readily seen in the weight profile of *W*^*rec*^ in Fig. 3C, and it is the substrate that allows the network to support stable activity bumps in virtually any location. In addition, we observe inhibition surrounding the local excitatory profile in both directions. This inhibition emerges despite the fact that we provide global inhibition to all HD cells (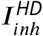 parameter, Methods), in line with suggestions from previous work (Kim et al., 2017). Surrounding inhibition was a feature we observed consistently in learned networks of different sizes and for different global inhibition levels. Finally, the angular offset of the two negative sidelobes in the connectivity depends on the size and shape of the entrained HD bump (for details, see Mathematical Appendix).

Furthermore, we find a consistent pattern of both L-HR and R-HR populations to excite the direction for which they are selective (Fig. 3C), which is also similar to what is observed in the CX (Turner-Evans et al., 2020). Excitation in one direction is accompanied by inhibition in the reverse direction in the learned network. As a result of the symmetry in our learning paradigm, the connectivity profiles of L-HR and R-HR cells are mirrored versions of each other, which is also clearly visible in Fig. 3C. The inhibition of the reverse direction has a width comparable to the bump size and acts as a “break” to prevent the bump from moving in this direction. The excitation in the selective direction, on the other hand, has a wider profile, which allows the network to path integrate for a wide range of angular velocities, i.e., for high angular velocities neurons further downstream can be “primed” and activated in rapid succession. Indeed, when we remove the wide projections from the excitatory connectivity, PI performance is impaired for the higher angular velocities exclusively (Fig. S1). The even weight profile in *W*^*rec*^ and the mirror symmetry for L-HR vs. R-HR profiles in *W*^*HR*^, together with the circular symmetry of the weights throughout the ring, guarantee that there is no side bias (i.e., tendency of the bump to favor one direction of movement versus the other) during PI. Indeed, the PI error distribution in Fig. 2B remains symmetric throughout the 60-s simulations.

Next, we focus our attention on the dynamics of learning. For training times larger than a few hours, the absolute learning error drops and settles to a low value, indicating that learning has converged after ∼20 hours of training time (Fig. 3D). The non-zero value of the final error is only due to errors occurring at the edges of the bump (Fig. S2A, top panel). An intuitive explanation of why these errors persist is that the velocity pathway is learning to predict the visual input; as a result, when the visual input is present, the velocity pathway creates errors that are consistent with PI velocity biases in darkness (see Supplementary Information).

Figures 3E,F show the weight development history for the entire simulation. The first structure that emerges during learning is the local excitatory recurrent connections in *W*^*rec*^. For these early stages of learning, the initial connectivity is controlled by the autocorrelation of the visual input, which gets imprinted in the recurrent connections by means of Hebbian co-activation of adjacent HD neurons. As a result, the width of the local excitatory profile mirrors the width of the visual input. Once a clear bump is established in the HD ring, the HR connections are learned to support bump movement, and negative sidelobes in *W*^*rec*^ emerge. To understand the shape of the learned connectivity profiles and the dynamics of their development, we study a reduced version of the full model, which follows learning in bump-centric coordinates (see Mathematical Appendix). The reduced model produces a connectivity strikingly similar to the full model, and highlights the important role of non-linearities in the system.

So far we have shown results in which our model far outperforms flies in terms of PI accuracy. To bridge this gap, we add noise to the weight connectivity in Fig. 3A,B, and obtain the connectivity matrices in Fig. S3A,B, respectively. This perturbation of the weights accounts for irregularities in the fly HD system owning to biological factors, such as uneven synaptic densities. The resulting neural velocity gain curve in Fig. S3E is impaired mainly for small angular velocities (cf. Fig. 2C). Interestingly, it now bears greater similarity to the one observed in flies, because the previously flat area for small angular velocities is wider (flat for *v <* 60 degrees/s, cf. extended data Fig. 7G,J in Seelig and Jayaraman (2015)). This happens because the noisy connectivity is less effective in initiating bump movement. Finally, the PI errors in the network with noisy connectivity grow much faster, and display a strong side bias (Fig. S3D, cf. Fig. 2B). The latter can be attributed to the fact that the noise in the connectivity generates local minima in the energy landscape of the ring attractor that are easier to transverse from one direction vs. the other. Side bias can also emerge if the learning rate *1* in eq. (15) is increased, effectively forcing learning to converge faster to a local minimum, which results in slight deviations from circularly symmetric connectivity (data not shown). It is therefore expected that different animals will display different degrees and directions of side bias during PI, owning either to fast learning or asymmetries in the underlying neurobiology. In the Supplementary Information we also incorporate random Gaussian noise to all inputs, which can account for noisy percepts or stochasticity of spiking, and show that learning is not disrupted even for high noise levels (Fig. S4).

### Fast adaptation to arbitrary neural velocity gains

Having shown how PI and CAN properties are learned in our model, we now turn our attention to the flexibility that our learning setup affords. Motivated by virtual-reality experiments in rodents where the relative gain of visual and self-motion inputs is manipulated (Jayakumar et al., 2019), we test whether our network can rewire to learn an arbitrary gain between the two. In other words, we attempt to learn an arbitrary gain *g* between the idiothetic angular velocity *v* sensed by the HR cells and the neural velocity *g v* dictated by the allothetic visual input. This simulates the conditions in a virtual reality environment, where the speed at which the world around the animal rotates is determined by the experimenter, but the proprioceptive sense of head angular velocity remains the same.

Starting with the learned network shown in Fig. 3, which displayed gain *g* = 1, we suddenly switch to a different gain, i.e., we learn weights for either *g* = 0.5 or *g* = 1.5, corresponding to a 50% decrease or increase in gain, respectively. In both cases, we observe that the network readily rewires to achieve the new gain. The mean learning error after the gain switch is initially high, but reaches a lower, constant level after only 1–2 hours of training (Fig. 4A). We note that convergence is much faster compared to the time it takes for the gain-1 network to emerge from scratch (compare to Fig. 3D). Importantly, Fig. 4B shows that PI performance in the resulting networks is excellent for the two new gains, with some degradation only for very low and very high angular velocities. There are two reasons why high angular velocities are not learned that well: limited training of these velocities, and saturation of HR cell activity. Both reasons are by design and do not reflect a fundamental limit of the network. In the Supplementary Information we show that without the aforementioned limitations the network learns to path-integrate up to an angular velocity limit set by synaptic delays (Fig. S5).

**Figure 4.**
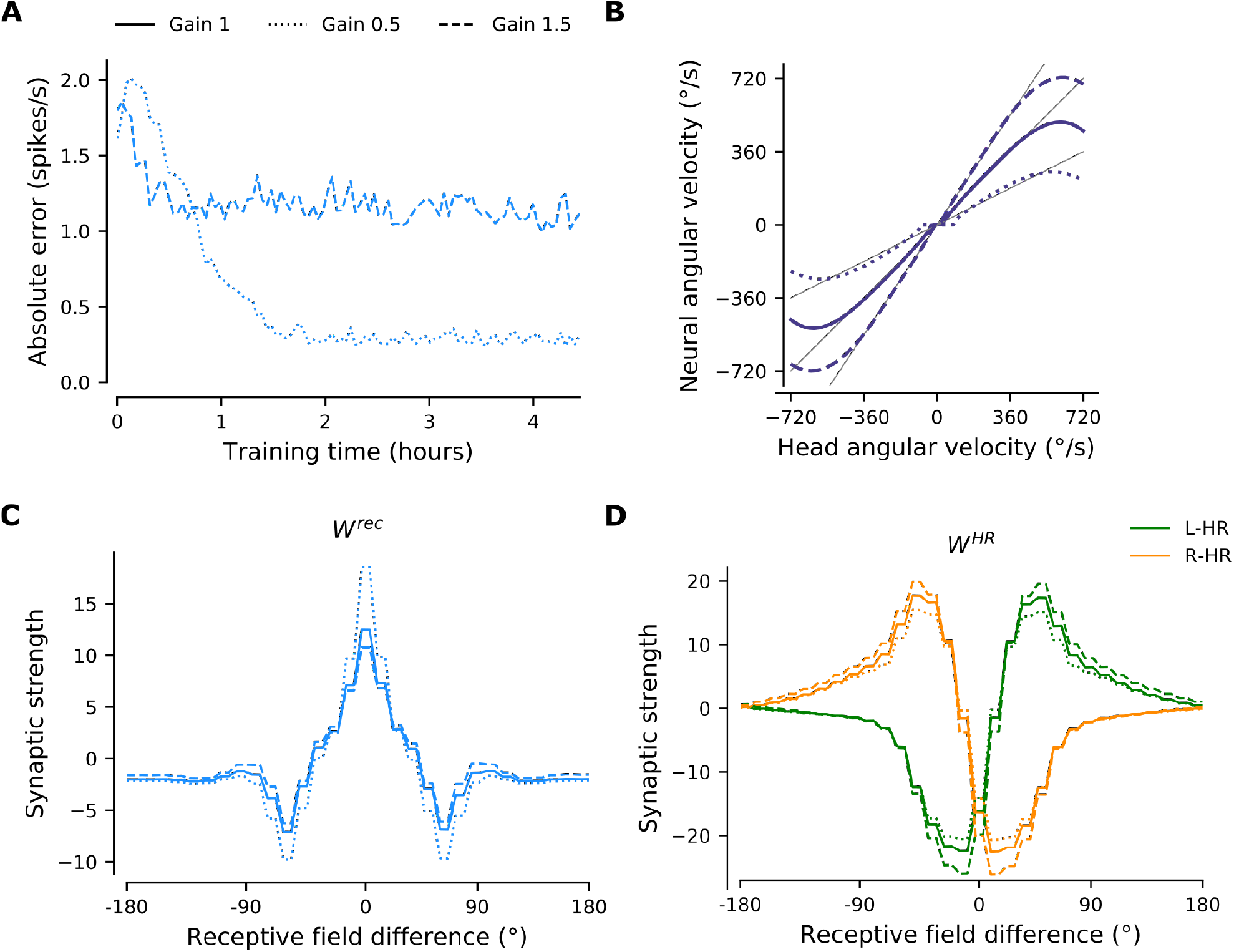
The network adapts rapidly to new gains. Starting from the converged network in Fig. 3, we change the gain between visual and self-motion inputs, akin to experiments conducted in VR in flies and rodents (Seelig and Jayaraman, 2015; Jayakumar et al., 2019). Solid, dotted, and dashed lines throughout the figure correspond to gains *g* = {1, 0.5, 1.5}, respectively. (A) The mean learning error averaged across 12 simulations for different gains. After an initial increase due to the change of gain, the errors decrease rapidly and settle to a low value. The steady-state values depend on the gain due to the by-design impairment of high angular velocities, which affects high gains preferentially. Crucially, adaptation to a new gain is much faster than learning the HD system from scratch (cf. Fig. 3D). (B) Velocity gain curves for different gains. The network has remapped to learn accurate PI with different gains for the entire dynamic range of head angular velocity inputs (approx. [−500, 500] deg/s). (C), (D) Final profiles of *W*^*rec*^ and *W*^*HR*^, respectively, for different gains.

Figures 4C,D compare the weight profiles of the resulting circularly symmetric matrices *W*^*rec*^ and *W*^*HR*^, respectively, for the three different gains. An increase in gain slightly suppresses the recurrent connections and slightly amplifies the HR-to-HD connections, while a decrease in gain substantially amplifies the recurrent connections and slightly suppresses the HR-to-HD connections. The latter explains why the flat region for small angular velocities in Fig. 4B has been extended for *g* = 0.5: it is now harder for small angular velocities to overcome the attractor formed by stronger recurrent weights and move the bump. We note that the network can also learn to reverse its gain (*g* = −1), i.e., when the visual and self-motion inputs are signaling movement in opposite directions. However, it takes considerably more time to do so than learning *g* = −1 from scratch (data not shown). Overall, our simulations demonstrate that the visual input exerts stronger control on plasticity in the network than the velocity pathway, leading to rewiring of the PI circuit.

## Discussion

The ability of animals to navigate in the absence of external cues is crucial for their survival. Head direction, place, and grid cells provide internal representations of space (Ranck, 1984; Moser et al., 2008) that can persist in darkness and possibly support path integration (PI) (Mizumori and Williams, 1993; Quirk et al., 1990; Hafting et al., 2005). Extensive theoretical work has focused on how the spatial navigation system might rely on continuous attractor networks (CANs) to maintain and update a neural representation of the animal’s current location. Special attention was devoted to models representing orientation, with the ring attractor network being one of the most famous of these models (Amari, 1977; Ben-Yishai et al., 1995; Skaggs et al., 1995; Seung, 1996). So far, modelling of the HD system has been relying on hand-tuned synaptic connectivity (Zhang, 1996; Xie et al., 2002; Turner-Evans et al., 2017; Page et al., 2018) without reference to its origin; or has been relying on synaptic plasticity rules that either did not achieve gain-1 PI (Stringer et al., 2002) or were not biologically plausible (Hahnloser, 2003).

### Summary of ndings

Inspired by the recent discovery of a ring attractor network for HD in *Drosophila* (Seelig and Jayaraman, 2015), we show how a biologically plausible learning rule leads to the emergence of a circuit that achieves gain-1 PI in darkness. The learned network features striking similarities in terms of connectivity to the one experimentally observed in the fly (Turner-Evans et al., 2020), and reproduces experiments on CAN dynamics (Kim et al., 2017) and gain changes between external and self-motion cues (Jayakumar et al., 2019). Furthermore, an emergent property of the mature network is an impairment of PI for small angular velocities, which is a feature that has been reported in experiments (Seelig and Jayaraman, 2015).

The mature circuit displays two properties characteristic of CANs: 1) it can support and actively maintain a local bump of activity at a virtual continuum of locations, and 2) it can move the bump across the network by integrating self-motion cues. Note that we did not explicitly train the network to achieve these CAN properties, but they rather emerged in a self-organized manner.

To achieve gain-1 PI performance, our network must attribute learning errors to the appropriate weights. The learning rule we adopt is a “delta-like” rule, with a learning error in eq. (17) that gates learning in the network, and a Hebbian component that comes in the form of the postsynaptic potential in eq. (11) and assigns credit to synapses that are active when errors are large. The learning rule leads to the emergence of both symmetric local connectivity between HD cells (which is required for bump maintenance and stability), and asymmetric connectivity from HR to HD cells (which is required for bump movement in darkness). The first happens because adjacent neurons are co-active due to correlated visual input; the second because only one HR population is predominantly active during rotation: the population that corresponds to the current rotation direction (for details, see also the Mathematical Appendix).

### Relation to experimental literature

Our work comes at a time at which the fly HD system receives a lot of attention (Seelig and Jayaraman, 2015; Turner-Evans et al., 2020; Kim et al., 2017, 2019; Fisher et al., 2019), and suggests a mechanism of how this circuit could self-organize during development. Synaptic plasticity has been shown to be important in this circuit for anchoring the visual input to the HD neurons when the animal is exposed to a new environment (Kim et al., 2019; Fisher et al., 2019). This has also been demonstrated in models of the mammalian HD system (Skaggs et al., 1995; Zhang, 1996; Song and Wang, 2005). Here we assume that an initial anchoring of the topographic visual input to the HD neurons with arbitrary offset with respect to external landmarks already exists prior to the development of the PI circuit; such an anchoring could even be prewired. In our model, it is sufficient that the visual-input tuning is local and topographically arranged. Once the PI circuit has developed, visual connections could be anchored to different environments, as shown by Kim et al. (2019) and Fisher et al. (2019). For the sake of simplicity and without loss of generality, we study the development of the path-integrating circuit while the animal moves in the same environment, and keep the visual input-tuning fixed. Therefore, the present work addresses the important question of how the PI circuit itself could be formed, and it is complementary to the problem of how allothetic inputs to the PI circuit are wired (Fisher et al., 2019; Kim et al., 2019).

In Figure 4 we show that our network can adapt to different gains much faster than the time required to learn the network from scratch. Our simulations are akin to experiments where rodents are placed in a VR environment and the relative gain between visual and proprioceptive signals is altered by the experimenter (Jayakumar et al., 2019). In this scenario, Jayakumar et al. (2019) found that the PI gain of place cells can be recalibrated rapidly. In contrast, Seelig and Jayaraman (2015) found that PI gain in darkness is not significantly affected when flies are exposed to different gains in light conditions. We note, however, that Seelig and Jayaraman (2015) tested mature animals (8–11 days old), whereas plasticity in the main HD network is presumably stronger in younger animals. Also note that the manipulation we use to address adaptation of PI to different gains differs from the one in (Kim et al., 2019) who used optogenetic stimulation of the HD network combined with rotation of the visual scene to trigger a remapping of the visual input to the HD cells in a Hebbian manner. The findings in Jayakumar et al. (2019) can only be reconciled by plasticity in the PI circuit, and not in the sensory inputs to the circuit.

In order to address the core mechanisms that underlie the emergence of a path integrating network, we use a model that is a simplified version of the biological circuit. For example, we did not model inhibitory neurons explicitly and omitted some of the recurrent connectivity in the circuit, whose functional role is uncertain (Turner-Evans et al., 2020). We also choose to separate PI from other complex processes that occur in the CX (Raccuglia et al., 2019). Finally, we do not force the network to obey Dale’s law and do not model spiking explicitly.

Nevertheless, after learning, we obtain a network connectivity that is strikingly similar to the one of the fly HD system. Indeed, the mature model exhibits local excitatory connectivity in the HD neurons, which in the fly is mediated by the excitatory loop from E-PG to P-EG to P-EN2 and back to E-PG (Turner-Evans et al., 2020), a feature that hand-tuned models of the fly HD system did not include (Turner-Evans et al., 2017). Furthermore, the HR neurons have excitatory projections towards the directions they are selective for, similar to P-EN1 neurons in the fly. Interestingly, these key features that we uncover from learning have been utilized in other hand-tuned models of the system (Turner-Evans et al., 2017; Kim et al., 2017, 2019). Future work could endeavor to come closer to the architecture of the fly HD system and benefit from the incorporation of more neuron types and the richness of recurrent connectivity that has been recently discovered in the fly (Turner-Evans et al., 2020).

### Relation to theoretical literature

A common problem with CANs is that they require fine tuning: even a slight deviation from the optimal synaptic weight tuning leads to catastrophic drifting (Goldman et al., 2009). A way around this problem is to sacrifice the continuity of the attractor states in favor of a discrete number of stable states that are much more robust to noise or weight perturbations (Kilpatrick et al., 2013). In our network, the small number of HD neurons enforces by design a coarsegrained representation of heading; the network is a CAN only in a quasi-continuous manner, and the number of discrete attractors corresponds to the number of HD neurons. This makes it harder to “jump” to adjacent attractors, since an energy barrier has to be overcome in the quasi-continuous case (Kilpatrick et al., 2013). Overall, the quasi-continuous nature of the attractor shields the internal representation of heading against the ever-present biological noise, which would otherwise lead to diffusion of the bump with time. The fact that the network can still path-integrate accurately with this coarse-grained representation of heading is remarkable.

Seminal theoretical work on ring attractors has proven that in order to achieve gain-1 PI, the asymmetric component of the network connectivity (corresponding here to *W*^*HR*^) needs to be proportional to the derivative of the symmetric component (corresponding to *W*^*rec*^) (Zhang, 1996). However, this result rests on the assumption that asymmetric and symmetric weight profiles are mediated by the same neuronal population, as in the double-ring architecture proposed by Xie et al. (2002) and Hahnloser (2003), but does not readily apply to the architecture of the fly HD system where HD and HR cells are separate. In our learned network we find that the HR weight profile is not proportional to the derivative of the recurrent weight profile, therefore this requirement is not necessary for gain-1 PI in our setting. Note that our learning setup can also learn gain-1 PI for a double-ring architecture, which additionally obeys Dale’s law (P. Vafidis (2019), *Learning of a path-integrating circuit* [Unpublished master’s thesis]. Technical University of Berlin.).

Our learning setup, inspired by Urbanczik and Senn (2014), is similar to the one in Guerguiev et al. (2017) in the sense that both involve compartmentalized neurons that receive “target” signals in a distinct compartment. It differs, however, in the algorithm and learning rule used. Guerguiev et al. (2017) use local gradient descent during a “target” phase, which is separate from a forward propagation phase, akin to forward/backward propagation stages in conventional deep learning. In contrast, we use a modified Hebbian rule, and in our model “forward” computation and learning happen at the same time; time multiplexing, whose origin in the brain is unclear, is not required. Our setting would be more akin to the one in Guerguiev et al. (2017) if an episode of PI in darkness would be required before an episode of learning in light conditions, which does not seem in line with the way animals naturally learn.

### Outlook

The present study adds to the growing literature of potential computational abilities of compartmentalized neurons (Poirazi et al., 2003; Guerguiev et al., 2017; Beniaguev et al., 2019; Gidon et al., 2020). The associative HD neuron used in this study is a coincidence detector, which serves to associate external and internal inputs arriving at different compartments of the cell. Coupled with memory-specific gating of internally generated inputs, coincidence detection has been suggested to be the fundamental mechanism that allows the mammalian cortex to form and update internal knowledge about external contingencies (Doron et al., 2020). This structured form of learning does not require engineered “hints” during training (e.g. see DePasquale et al. (2018)), and it might be the reason why neural circuits evolved to be so efficient at reasoning about the world, with the mammalian cortex being the pinnacle of this achievement. Here we demonstrate that learning at the cellular level can predict external inputs (visual information) by associating firing activity with internally generated signals (velocity inputs) during training. This effect is due to the anti-Hebbian component of the learning rule in eq. (11), where the product of postsynaptic axon-distal and presynaptic activity comes with a negative sign. Specifically, it has previously been demonstrated that anti-Hebbian synaptic plasticity can stabilize persistent activity (Xie and Seung, 2000) and perform predictive coding (Bell et al., 1997; Hahnloser, 2003). At the population level, this provides a powerful mechanism to internally produce activity patterns that are identical to the ones induced from an external stimulus. This mechanism can serve as a way to anticipate external events or, as in our case, as a way of “filling in” missing information in the absence of external inputs.

Local, Hebb-like learning rules have been deemed a weak form of learning, due to their inability to utilize error information in a sophisticated manner. Despite that, we show that local associative learning can be particularly successful in learning appropriate fine-tuned synaptic connectivity, when operating within a cell structured for coincidence detection. Therefore, in learning and reasoning about the environment, our study highlights the importance of inductive biases with developmental origin (e.g. allothetic and idiothetic inputs arrive in different compartments of associative neurons), rather than powerful learning algorithms (Lake et al., 2016).

Importantly, our model generates testable predictions for future experiments in flies and, potentially, other animal models. An obvious prediction of our model is that synaptic plasticity is critical for the development of the PI network for heading, and the lack of allothetic sensory input (e.g. visual) during development will disrupt the formation of the PI system. Previous work showed that head direction cells in rat pups displayed mature properties already in their first exploration of the environment outside their nest (Langston et al., 2010), which may seem to contradict our assumption that the PI circuit wires during development. However, directional selectivity of HD cells in the absence of allothetic inputs and PI performance were not tested in this study. We also predict that HD neurons have a compartmental structure where idiothetic inputs are separated from allothetic sensory inputs, which initiate action potentials more readily due to being electrotonically closer to the axon initial segment. Finally, similarly to place cell studies in rodents (Jayakumar et al., 2019), we predict that during development the PI system can adapt to experimenter-defined gain manipulations, and it can do so faster than the time required for the system to develop from scratch.

In conclusion, the present work addresses the age-old question of how to develop a CAN that performs accurate, gain-1 PI in the absence of external sensory cues. We show that this feat can be achieved in a network model of the HD system by means of a biologically plausible learning rule at the cellular level. Even though our network architecture is tailored to the one of the fly CX, the learning setup is general and can be applied to other PI circuits. Of particular interest is the rodent HD system: despite the lack of evidence for a topographically-organized recurrent HD network in rodents, a one-dimensional HD manifold has been extracted in an unsupervised way (Chaudhuri et al., 2019). Therefore, our work lays the path to study the development of ring-like neural manifolds in mammals. Finally, it would be interesting to see if a similar mechanism underlies the emergence of PI in place and grid cells.

## Acknowledgements

We would like to thank Raquel Suárez-Grimalt and Marcel Heim for fruitful discussions. This work was funded by the Deutsche Forschungsgemeinschaft (DFG, German Research Foundation; SFB 1315 – project-ID 327654276 to R.K. and D.O.; and the Emmy Noether Programme 282979116 to D.O.), the German Federal Ministry for Education and Research (BMBF; Grant 01GQ1705 to R.K.), and the Onassis Foundation (P.V.).

## Author Contributions

P.V., T.D., and R.K. conceived the study. P.V. performed analyses and wrote the initial draft of the manuscript. T.D. contributed to analyses. D.O., T.D., and R.K. supervised the research. All authors wrote the manuscript.

## Methods

In what follows, we describe our computational model for learning a ring attractor network that accomplishes accurate angular PI. The model described here focuses on the HD system of the fly, however the proposed computational setup is general and could be applied to other systems. All parameter values are summarized in Table 1.

**Table 1.**
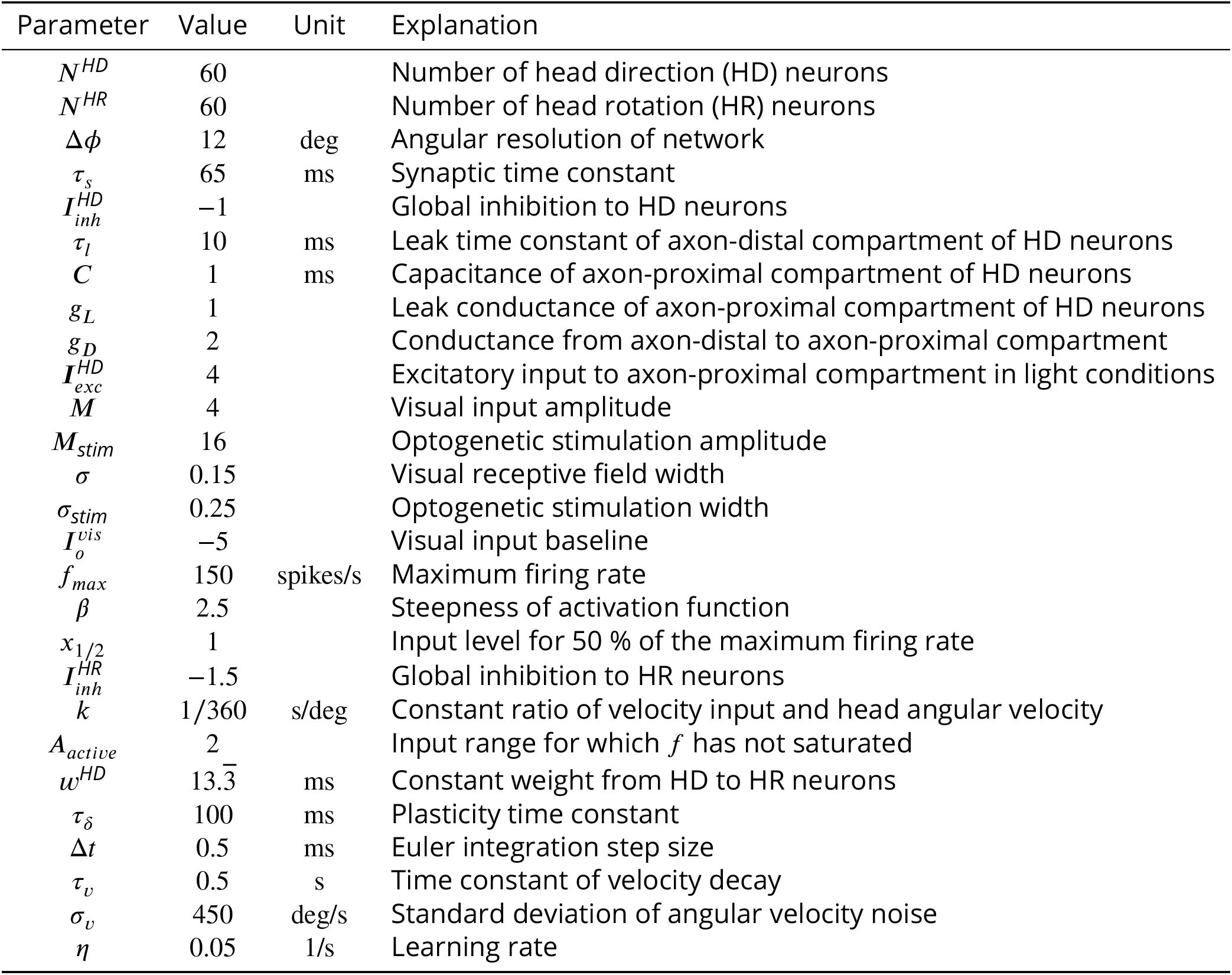
Parameter values, in the order they appear in the Methods section. These values apply to all simulations, unless otherwise stated. Note that voltages, currents, and conductances are assumed unitless in the text; therefore capacitances have the same units as time constants.

## Network Architecture

We model a recurrent neural network comprising *N*^*HD*^ = 60 head-direction (HD) and *N*^*HR*^ = 60 head-rotation (HR) cells, which are close to the number of E-PG and P-EN1 cells in the fly central complex (CX), respectively (Turner-Evans et al., 2020; Xu et al., 2020). A scaled-down version of the network for *N*^*HR*^ = *N*^*HD*^ = 12 is shown in Fig. 1A. The average spiking activity of HD and HR cells is modelled by firing-rate neurons. HD cells are organized in a ring and receive visual input, which encodes the angular position of the animal’s head with respect to external landmarks. We use a discrete representation of angles and we model two HD cells for each head direction, as observed in the biological system (Turner-Evans et al., 2017). Therefore the network can represent head direction with an angular resolution Δ*Ø* = 12 deg.

Motivated by the anatomy of the fly CX (Green et al., 2017; Turner-Evans et al., 2020), HR cells are divided in two populations (Fig. 1A): a ‘leftward’ (L-HR) population (with increased velocity input when the head turns leftwards) and a ‘rightward’ (R-HR) population (with increased velocity input when the head turns rightwards). After learning, these two HR populations will be responsible to move the HD bump in the anticlockwise and clockwise directions, respectively.

The recurrent connections among HD cells and the connections from HR to HD cells are assumed to be plastic. On the contrary, connections from HD to HR cells are assumed fixed and determined as follows: for every head direction, one HD neuron projects to a cell in the L-HR population, and the other to a cell in the R-HR population. Because HD cells project to HR cells in a 1-to-1 manner, each HR neuron is simultaneously tuned to a particular head direction and a particular head rotation direction. The synaptic strength of the HD to HR projections is the same for all projections. Finally, HR cells do not form recurrent connections.

### Neuronal Model

We assume that each HD neuron is a rate-based associative neuron (Fig. 1D), i.e., a two-compartmental neuron comprising an axon-proximal and an axon-distal dendritic compartment (Urbanczik and Senn, 2014; Brea et al., 2016). The two compartments model the dendrites of that neuron that are closer to (further away from) the axon initial segment. Note that here the axon-proximal compartment replaces the somatic compartment in the original model by Ur- banczik and Senn (2014). This is because the somata of fly neurons are typically electrotonically segregated from the rest of the cell and they are assumed to contribute little in computation (Gouwens and Wilson, 2009; Tuthill, 2009). We also note that to fully capture the input/output transformations that HD neurons in the fly perform, more compartments than two might be needed (Xu et al., 2020). Finally, only HD cells are associative neurons, whereas HR cells are simple rate-based point neurons.

HD cells receive an input current ***I***^*d*^ to the axon-distal dendrites, which obeys

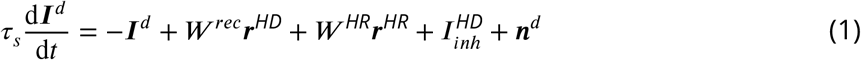

where ***I***^*d*^ is a vector of length *N*^*HD*^ with each entry corresponding to one HD cell. In eq. (1), *r*_*s*_ is the synaptic time constant, *W*^*rec*^ is a *N*^*HD*^ × *N*^*HD*^ matrix of the recurrent synaptic weights among HD cells, *W*^*HR*^ is a *N*^*HD*^ × *N*^*HR*^ matrix of the synaptic weights from HR to HD cells, ***r***^*HR*^ and ***r***^*HD*^ are vectors of the firing rates of HR and HD cells respectively, 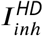 is a constant inhibitory input common to all HD cells, and ***n***^*d*^ are random fluctuations in the axon-distal input (noise) drawn IID from a Gaussian distribution with zero mean and variance. 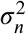. Note that in the main text we set *σ*_*n*_ to zero, but we explore different values for this parameter in the Supplementary Information (see section “Robustness to noise”). The constant current 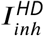 is in line with a global inhibition model with local recurrent connectivity, as opposed to having long-range inhibitory recurrent connectivity (Kim et al., 2017). The inhibitory current 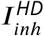 contributes to preserve the uniqueness of the HD bump, however the exact strength of this inhibition is not important in our model.

Since several electrophysiological parameters of the fly neurons modeled here are unknown, we use dimensionless conductance values. Therefore, in eq. (1), which describes the dynamics of the axon-distal input of HD cells, currents (e.g., ***I***^*d*^, 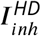, and ***n***^*d*^) are dimensionless. As a result, voltages are also dimensionless, and because we measure firing rates in units of 1/s, all synaptic weights (e.g., *W*^*rec*^ and *W*^*HR*^) then have, strictly speaking, the unit ‘seconds’ (s), even though we mostly suppress this unit in the text. Importantly, all time constants (e.g., *τ*_*s*_), which define the time scale of dynamics, are measured in units of time (in seconds).

The axon-distal voltage ***V***^*d*^ of HD cells is a low-pass filtered version of the input current ***I***^*d*^, that is,

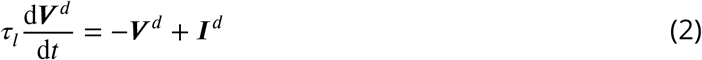

where *τ*_*l*_ is the leak time constant of the axon-distal compartment. The voltage ***V***^*d*^ and the current ***I***^*d*^ having the same unit (both dimensionless) means that the leak resistance of the axon-distal compartment is also dimensionless, and we assume that it is unity for simplicity. We choose values of *τ*_*l*_ and *τ*_*s*_ (for specific values, see Table 1) so that their sum matches the phenomenological time constant of HD neurons (E-PG in the fly), while *τ*_*s*_ equals to the phenomenological time constant of HR neurons (P-EN1 in the fly, Turner-Evans et al., 2017).

The axon-proximal voltage ***V*** ^***a***^ of HD cells is then given by

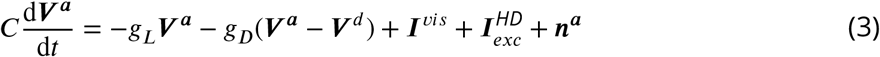

where *C* is the capacitance of the membrane of the axon-proximal compartment, *g*_*L*_ is the leak conductance, *g*_*D*_ is the conductance of the coupling from axon-distal to axon-proximal dendrites, ***I***^*vis*^ is a vector of visual input currents to the axon-proximal compartment of HD cells, 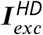 is an excitatory input to the axon-proximal compartment, and ***n***^***a***^ is a vector of IID Gaussian noise with zero mean and variance 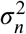 injected to the axon-proximal compartment. The excitatory current 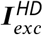, is assumed to be present only in light conditions. The values of *C, g*_*L*_, and *g*_*D*_ in the fly HD (E-PG) neurons are unknown, thus we keep these parameters unitless, and set their values to the ones in Urbanczik and Senn (2014). Note that since conductances are dimensionless here, *C* is effectively a time constant.

Following Hahnloser (2003), the visual input to HD cell *i* is a localized bump of activity at angular location *θ*_*i*_:

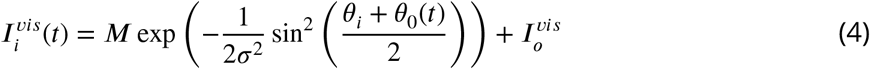

where *M* scales the bump’s amplitude, *σ* controls the width of the bump, *θ*_*i*_ is the preferred orientation of HD neuron *i, θ*_0_(*t*) is the position of a visual landmark at time *t* in head-centered coordinates, and 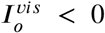 is a constant inhibitory current that acts as the baseline for the visual input. We choose *M* so that the visual input can induce a weak bump in the network at the beginning of learning, and we choose *σ* so that the resulting bump after learning is ∼60 degrees wide. Note that the bump in the mature network has a square shape (Fig. S2B); therefore we elect to make it slightly narrower than the average full width at half maximum of the experimentally observed bump (∼80 degrees; Seelig and Jayaraman, 2015; Kim et al., 2017; Turner-Evans et al., 2017). In addition, the current 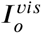 is negative enough to make the visual input purely inhibitory, as reported (Fisher et al., 2019). The visual input is more inhibitory in the surround to suppress activity outside of the HD receptive field. Therefore the mechanism in which the visual input acts on the HD neurons is disinhibition.

The firing rate of HD cells, which is set by the voltage in the axon-proximal compartment, is given by

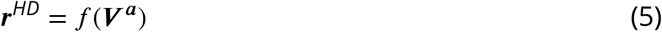

where

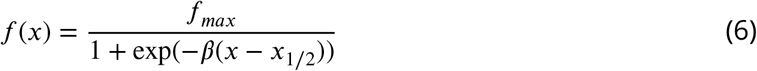

is a sigmoidal activation function applied element-wise to the vector ***V***. The variable *f*_*max*_ sets the maximum firing rate of the neuron, *β* is the slope of the activation function, and *x*_1/2_ is the input level at which half of the maximum firing rate is attained. The value of *f*_*max*_ is arbitrary, while *β* is chosen such that the activation function has sufficient dynamic range and *x*_1/2_ is chosen such that for small negative inputs the activation function is non-zero.

The firing rates of the HR cells are given by

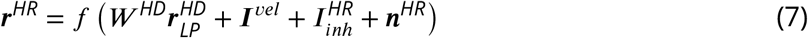

where ***r***^*HR*^ is the vector of length *N*^*HR*^ of firing rates of HR cells, the *N*^*HR*^ × *N*^*HD*^ matrix *W*^*HD*^ encodes the fixed connections from the HD to the HR cells, 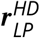 is a low-pass filtered version of the firing rate of the HD cells where the filter accounts for delays due to synaptic transmission in the incoming synapses from HD cells, ***I***^*vel*^ is the angular velocity input, 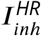 is a constant inhibitory input common to all HR cells, and ***n***^*HR*^ is a IID Gaussian noise input to the HR cells with zero mean and variance 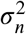. We set 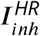 to a value that still allows sufficient activity in the HR cells bump, even when the animal does not move. The low-pass filtered firing-rate vector 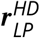 is given by

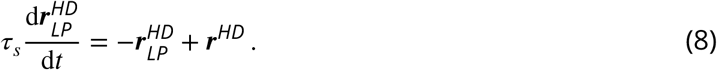

and the angular-velocity input to HR neuron *i* is given by

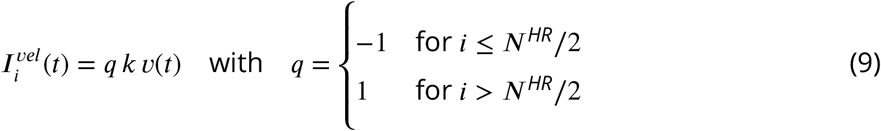

where *k* is the proportionality constant between head angular velocity and velocity input to the network, *v*(*t*) is the head angular velocity at time *t* in units of deg/s, and the factor *q* is chosen such that the left (right) half of the HR cells are primarily active during leftward (rightward) head rotation. Note that the same *τ*_*s*_ is in both eqs. (1) and (8). Finally, as mentioned earlier, the matrix *W*^*HD*^ encodes the hardwired HD-to-HR connections, i.e., 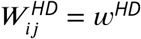 if HD neuron *j* projects to HR neuron *i*, and 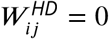 otherwise. Specifically, for *j* odd, HD neuron projects to L-HR neuron 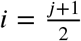, whereas for *j* even, HD neuron projects to R-HR neuron 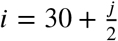. The synaptic strength *w*^*HD*^ is chosen such that the range of the firing rates of the HD cells is mapped to the entire range of firing rates of the HR cells. Specifically, we set 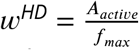, where *A*_*active*_ is the range of inputs for which *f* has not saturated, i.e., the input values for which *f* remains between about 7% and 93% of its maximum firing rate *f*_*max*_ (see eq. (6)). Finally, the proportionality constant *k* is set so that the firing rate of HR neurons does not reach saturation for the range of velocities relevant for the fly (approx. [−500, 500] deg/s), given all other inputs they receive.

### Synaptic Plasticity Rule

In our network, the associative HD neurons receive direct visual input in the axon-proximal compartment and indirect angular velocity input in the axon-distal compartment through the HR-to-HD connections (Fig. 1D). We hypothesize that the visual input acts as a supervisory signal that controls the axon-proximal voltage ***V*** ^***a***^ directly, and the latter initiates spikes. Therefore, the goal of learning is for the axon-distal voltage ***V*** ^*d*^ to predict the axon-proximal voltage by changing the synaptic weights *W*^*rec*^ and *W*^*HR*^. This change is achieved by minimizing the difference between the firing rate *f* (***V*** ^***a***^) in the presence of visual input and the axon-distal prediction *f* (***V*** ^*ss*^) of the firing rate in the absence of visual input. In the latter case and at steady-state, the voltage 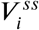 for HD neuron *i* is an attenuated version of the axon-distal volt-age,

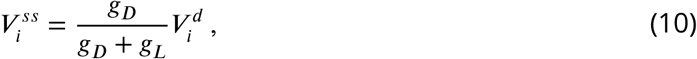

with conductance *g*_*D*_ of the coupling from the axon-distal to axon-proximal dendrites and leak conductance *g*_*L*_ of the axon-proximal compartment, as explained in eq. (3). Therefore, following Urbanczik and Senn (2014), we define the plasticity-induction variable *Pl*_*ij*_ for the connection between presynaptic neuron *j* and postsynaptic neuron *i* as

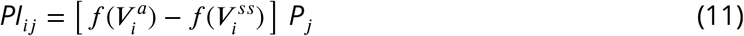

where *P*_*j*_ is the postsynaptic potential of neuron *j* which is a low-pass filtered version of the presynaptic firing rate *r*_*j*_ That is,.

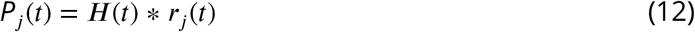

where * denotes convolution. The transfer function

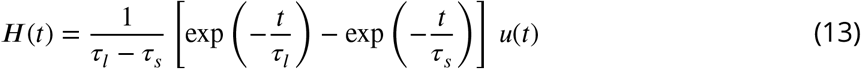

is derived from the filtering dynamics in eq. (1) and eq. (2) and accounts for the delays introduced by the synaptic time constant *τ*_*s*_ and the leak time constant *τ*_*l*_. In eq. (13), *u*(*t*) denotes the Heaviside step function, i.e., *u*(*t*) = 1 for *t >* 0 and *u*(*t*) = 0 otherwise. The plasticity-induction variable is then low-pass filtered to account for slow learning dynamics,

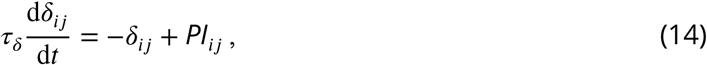

and the final weight change is given by

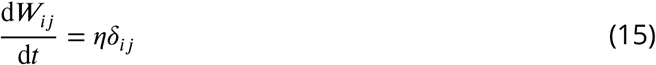

where *η* is the learning rate and *W*_*ij*_ is the connection weight from presynaptic neuron *j* to postsynaptic neuron *i*. Note that the synaptic weight *W*_*ij*_ is an element of either the matrix *W*^*rec*^ or the matrix *W*^*HR*^ depending on whether the presynaptic neuron *j* is an HD or an HR neuron, respectively. The value of the plasticity time constant *τ*_*δ*_ is not known, therefore we adopt the value suggested by Urbanczik and Senn (2014).

Equation (11) is a “delta-like” rule that can be interpreted as an extension of the Hebbian rule; compared to a generic Hebbian rule, we have replaced the postsynaptic firing rate 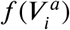 by the difference between 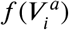 and the predicted firing rate 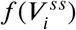 of the axon-distal compartment of the postsynaptic neuron. This difference drives plasticity in the model. We note that 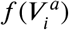 is a continuous approximation of the spike train of the postsynaptic neuron, which could be available at the axon-distal compartment via back-propagating action potentials (Larkum, 2013). Furthermore, the axon-distal voltage and postsynaptic potentials are by definition available at the synapses arriving at the axon-distal compartment. Therefore, the learning rule is biologically plausible because all information is locally available at the synapse.

The learning rule used here differs from the one in the original work of Urbanczik and Senn (2014) because we utilize a rate-based version instead of the original spike-based version. Even though spike trains can introduce Poisson noise to 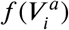, Urbanczik and Senn (2014) show that once learning has converged, asymmetries in the weights due the spiking noise are on average canceled out.

Another difference in our learning setup is that, unlike in Urbanczik and Senn (2014), the input to the axon-proximal compartment does not reach zero in equilibrium (see, e.g., Fig. 3D, and the Mathematical Appendix). Therefore, an activation function with a saturating non-linearity, as in eq. (6), is crucial for convergence, which could not be achieved with a less biologically plausible threshold-linear activation function. This lack of strict convergence in our setup is responsible for the square form of the bump (Fig. S3B and Mathematical Appendix).

### Training Protocol

We train the network with synthetically generated angular velocities, simulating head turns of the animal. The network dynamics are updated in discrete time steps Δ.*t* using forward Euler integration. The entrained angular velocities cover the range of angular velocities exhibited by the fly, which are at maximum ∼ 500 deg/s during walking or flying (Geurten et al., 2014; Stowers et al., 2017). The angular velocity *v*(*t*) is modeled as an Ornstein–Uhlenbeck process given by

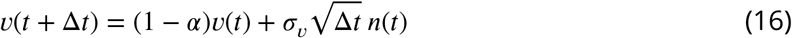

where *α* = Δ.*t*/ *τ* _*v*_ and *τ*_*v*_ is the time constant with which *v*(*t*) decays to zero, *n*(*t*) is noise drawn from a normal distribution with mean 0 and standard deviation 1 at each time step, and *σ*_*v*_ scales the noise strength.

We pick *σ*_*v*_ and *τ*_*v*_ so that the resulting angular velocity distribution in Fig. S2C and its time course, e.g. in Fig. 2A, are similar to what has been reported in flies during walking or flying (Geurten et al., 2014; Stowers et al., 2017). Finally, note that we train the network for angular velocities a little larger than what flies typically display (up to ±720 deg/s).

### Quanti cation of the Mean Learning Error

In eq. (11) we have used the learning error

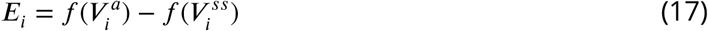

which controls learning in every associative HD neuron *i*. To quantify the mean learning error *err*(*t*) in the whole network at time *t*, we average *E*_*i*_ across all HD neurons and across a small time interval [*t, t* + *t*_*w*_], that is,

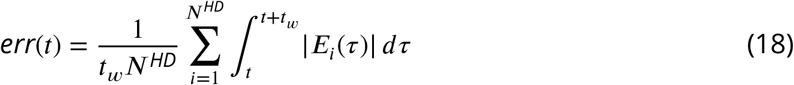

with *t*_*w*_ = 10 s. In Fig. 3D, we plot this mean error at every 1 % of the simulation, for 12 simulations, and averaged across the ensemble of the simulations. Note that individual simulations occasionally display “spikes” in the error. Large errors occur if the network happens to be driven by very high velocities that the network does not learn very well because they are rare; larger errors also occur for very small velocities, i.e., when the velocity input is not strong enough to overcome the local attractor dynamics, as seen, e.g., in Fig. 2C. On average, though, we can clearly see that the mean learning error decreases with increasing time and settles to a small value (e.g., Fig. 3D and Fig. 4A).

### Population Vector Average

To decode from the activity of HD neurons an average HD encoded by the network, we use the population vector average (PVA). We thus first convert the tuning direction *θ*_*i*_ of each HD neuron *i* to the corresponding complex number 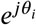 on the unitary circle, where is the imaginary unit. This complex number is multiplied by the firing rate 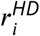 of HD neuron *i*, and then averaged across neurons to yield the PVA

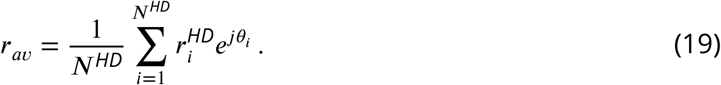

The PVA is a vector in the 2-D complex plane and points to the center of mass of activity in the HD network. Finally, we take the angle of the PVA as a measure for the current heading direction represented by the network.

### Fly Connectome Analysis

Our model assumes the segregation of visual inputs to HD (E-PG) cells from head rotation and recurrent inputs to the same cells. To test this hypothesis, we leverage on the recently released fly hemibrain connectome (Xu et al., 2020; Clements et al., 2020). First, we randomly choose one E-PG neuron per wedge of the EB, for a total of 16 E-PG neurons. We then find all incoming connections to these neurons from visually responsive ring neurons R2 and R4d (Omoto et al., 2017; Fisher et al., 2019). These are the connections that arrive at the axon-proximal compartment in our model. We then find all incoming connections from P-EN1 cells, which correspond to the HR neurons, and from P-EN2 cells, which are involved in a recurrent excitatory loop from E-PG to P-EG to P-EN2 and back to E-PG (Turner-Evans et al., 2020). These are the connections that arrive at the axon-distal compartment in our model.

## Supplementary Information

### Details of Learning

As further evidence that training in Fig. 3 of the main text has converged, we plot in Fig. S2A the learning error (eq. (17)) of the HD cells for the first 30 s of the same example simulation as Fig. 2A, and for both conditions. In light conditions, the error is zero in all positions apart from the edges of the bump, where the error is substantial. There exists an intuitive explanation of why these errors persist. During learning, the circuit is trying to predict the visual input, otherwise the additional delays in the velocity pathway (due to the extra synaptic transmission to HR neurons) would make it impossible to follow changes in the visual input perfectly. Moreover, the velocity pathway, which implements PI, cannot move the bump for very small angular velocities, and tends to move it slightly faster for intermediate velocities, and slower for large ones. Both of the latter biases are likely due to the saturation of angular velocities that the circuit integrates. Since the velocity pathway is active even in the presence of visual input, it creates errors at the edges of the bump whose sign is consistent with the aforementioned PI velocity biases (Fig. S2A, top panel). Other than that, the angular velocity input predicts the visual input near-perfectly, as evidenced by the near-zero error everywhere else in the network. Therefore, this strongly argues that learning has converged. During PI in darkness, the network operates in a self-consistent manner, merely integrating the angular velocity input, and the learning error is very small. Note that errors are not defined for HR cells since they are not associative neurons.

In addition, it is clear from Fig. S2B that the bump in the network has a square form, in contrast to the smoother form which would be expected from visual input alone. This is because the learning rule in eq. (11) only converges when HD neurons reach saturation (see also Mathematical Appendix, Fig. S8 panel S2).

### Robustness to noise

Up to this point, the only source of stochasticity in the network came from the angular velocity noise in the Ornstein-Uhlenbeck process (Methods). Biological HD systems, however, are subject to other forms of biological noise, like noisy percepts or stochasticity of spiking. To address that, we include Gaussian IID synaptic noise to every location in the network where inputs arrive: the axon-proximal and axon-distal compartments of HD cells and the HR cells (see Methods). We then ask how robustly can the network learn in the presence of such stochasticity.

To quantify the network’s robustness to noise, we need to define a comparative measure of useful signals vs. noise in the network. By “signals” we refer to the velocity/visual inputs and any network activity resulting from them, whereas “noise” is the aforementioned Gaussian IID variables. We then define the signal-to-noise ratio (*SNR*) to be the squared ratio of the active range of the activation function *f, A*_*active*_, over two times the standard deviation of the Gaussian noise, *σ*_*n*_, i.e.

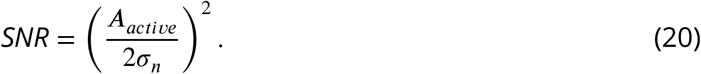

This definition is motivated by the fact that *A*_*active*_ determines the useful range signals in the network can have. If any of the signals exceed this range, they cannot impact the network in any meaningful way because the neuronal firing rate has saturated, unless they are counterbalanced by other signals reliably present. The 2 in the denominator is due to the fact that the noise can extend to both positive and negative values, whereas *A*_*active*_ denotes the entire range of useful inputs.

We then vary the *SNR* systematically and observe its impact on learning and network performance. Figure S5 shows a network that has been trained with *SNR* = 2. The resulting network connectivity remains circularly symmetric and maintains the required asymmetry in the HR-to-HD connections for L- and R-HR cells (data not shown). Therefore we only plot the profiles in Fig. S4B, which look very similar to the one in Fig. 3C. The network activity still displays a clear bump that smoothly follows the ground truth in the absence of visual input (Fig. S4C). There are only minor differences compared to the network without noise. The presence of the noise is most obvious in the HR cells, since HD cells that do not participate in the bump are deep into inhibition, and therefore synaptic input noise does not affect as much their activity. Also, we observe that the peak of the local excitatory connectivity in *W*^*rec*^ is not as pronounced. This happens because the noise corrupts auto-correlations of firing during learning. Further-more, the PI errors diffuse faster in the network with noise (Fig. S4D), and the neural velocity slightly overestimates the head angular velocity (Fig. S4E). In total, however, we conclude that the learning dynamics average out the impact of input noise, and the resulting network is excellent at PI, even when the *SNR* is low. Networks with higher *SNR* performed even better, whereas the network can no longer sustain a bump in darkness when *SNR* = 1, i.e. when the standard deviation of the noise covers the full active range of inputs (data not shown).

### Delays in the network set a limit for neural velocity during path integration

In the main text we trained the network for a set of angular velocities that cover the full range exhibited by the fly, and we showed that it can account for several key experimental findings. However, the ability of any continuous attractor network to path-integrate is naturally limited for high angular velocities, due to the synaptic delays inherent in any such network (Zhong et al., 2020). To evaluate the ability of our network to integrate angular velocities, we sought to identify a limit of what velocities could be learned.

The width of the HD bump in our network in degrees is here termed *BW*, and it is largely determined by the width *σ* of the visual receptive field. This is because during training we force the network to produce a bump with a width matching that of the visual input, when the latter is not present. In turn, the width of the learned local excitatory connectivity profile in *W* ^*rec*^ that guarantees such stable bumps of activity will be similar to the width of the bump, because recurrent connections during learning are only drawn from active neurons (non-zero *P*_*j*_ in eq. (11)). As mentioned in the main text, this emphasizes the Hebbian component of our learning rule (fire together - wire together). As a result, the width of local excitatory recurrent connections should be approximately *BW* degrees.

We reason that this width determines how fast the bump can move in the HD network. On first thought this seems counter-intuitive, because the HR neurons are responsible for moving the bump. However as we show in Fig. S1 the higher velocities are served by the long range excitatory connections in *W*^*HR*^, which are not strong enough to move the bump by themselves. Therefore contribution from HD cells is still needed, at least for high angular velocities. Then how fast this contribution can happen is limited by the delays in the network, since any self-motion or recurrent input must pass through the synaptic delays of HD neurons before it can impact the current head direction. Therefore, assuming that only one hop downstream is needed to move the bump to the next position, we reason that the maximum velocity that the network can achieve without external guidance (i.e. without visual input) is inversely proportional to the total one-hop delays in the HD network *τ*_*tot*_ ≈ *τ*_*s*_ + *τ*_*l*_, i.e.

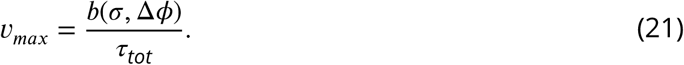

where *b* can be thought of as an effective bump width, which depends on *σ* but also on the angular resolution of the HD network Δ*ϕ*, due to discretization effects.

We then systematically vary the synaptic delays in the network and test what velocities it can learn. We indeed find that networks can path-integrate all angular velocities up to a limit, but not higher than that. As predicted, this limit is inversely proportional to the total one-hop delays in the HD network *τ*_*tot*_, for a wide range of delays (Fig. S5A). Furthermore, the inverse proportionality constant matches *BW* well. Fitting eq. (21) to the data we obtain *b*(0.25, 6) ≈ *BW* = 96 deg for *N*^*HD*^ = *N*^*HR*^ = 120 and *b*(0.15, 12) = 80 deg, *BW* = 60 deg for *N*^*HD*^ = *N*^*HR*^ = 60. The velocity gain plot for an example network with high synaptic delays is shown in Fig. S5B. Interestingly, we notice that the performance drop at the velocity limit is not gradual; instead, the neural velocity abruptly drops to a near-zero value once past the velocity limit. Further investigation reveals that for velocities higher than this limit, the network can no longer sustain a bump (Fig. S5C). This happens because the HD network cannot activate neurons downstream fast enough to keep the bump propagating, and therefore the bump disappears and the velocity gain plot becomes flat.

As mentioned in the main text, there are two limitations other than synaptic delays why the network could not learn high angular velocities: limited training of these velocities, and saturation of HR cell activity. These limitations kick in for total synaptic delays smaller than 160 ms. Therefore to create Fig. S5A for these delays, we increased the standard deviation of the velocity noise in the Ornstein–Uhlenbeck process to *σ*_*v*_ = 800 deg/s to address the first limitation, and we increased the dynamic range of angular velocity inputs by decreasing the proportionality constant to *k* = 1/540 s/deg to address the second.

Overall, these results indicate that the network learns to path-integrate all angular velocities up to a fundamental limit imposed by the architecture of the HD system in the fly. Furthermore, we conclude that the phenomenological delays observed in the fly HD system in Turner-Evans et al. (2017) are not fundamentally limiting the system’s performance, since they can support PI for angular velocities much higher than the ones normally displayed by the fly. Finally, the findings suggest that there is a trade-off between the bump width *BW* and the maximum angular velocity the HD system can support. Ideally, an animal would benefit from a small *BW* because it translates to a finer internal estimate of heading. However, since *b* in eq. (21) depends on *BW*, a smaller *BW* leads to reduced maximum angular velocity. Therefore, there exists a fundamental trade-off between location and velocity accuracy in the HD system.

## Supplementary Figures

**Figure S1.**
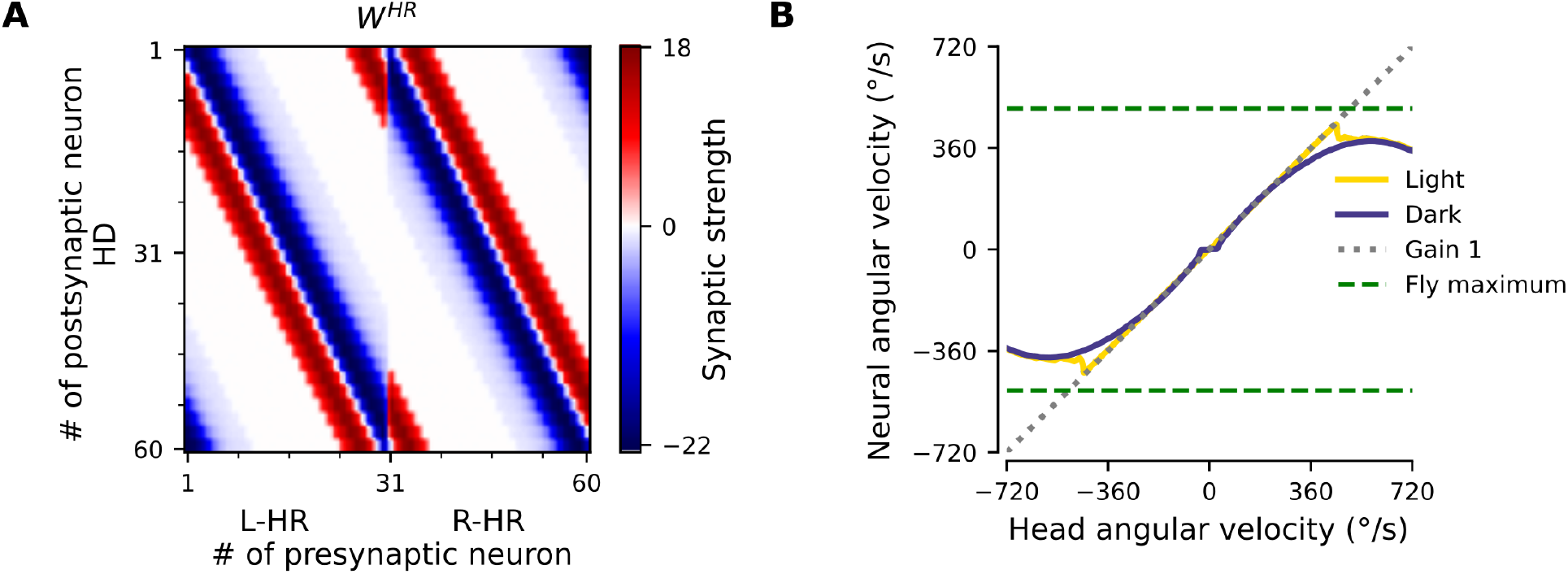
Removal of long-range excitatory projections impairs PI for high angular velocities. (A) The weight matrix *W*^*HR*^ of connections from HR to HD neurons from Fig. 3B, after the long-range excitatory projections have been removed. (B) PI in the resulting network is impaired for high angular velocities, compared to Fig. 2C.

**Figure S2.**
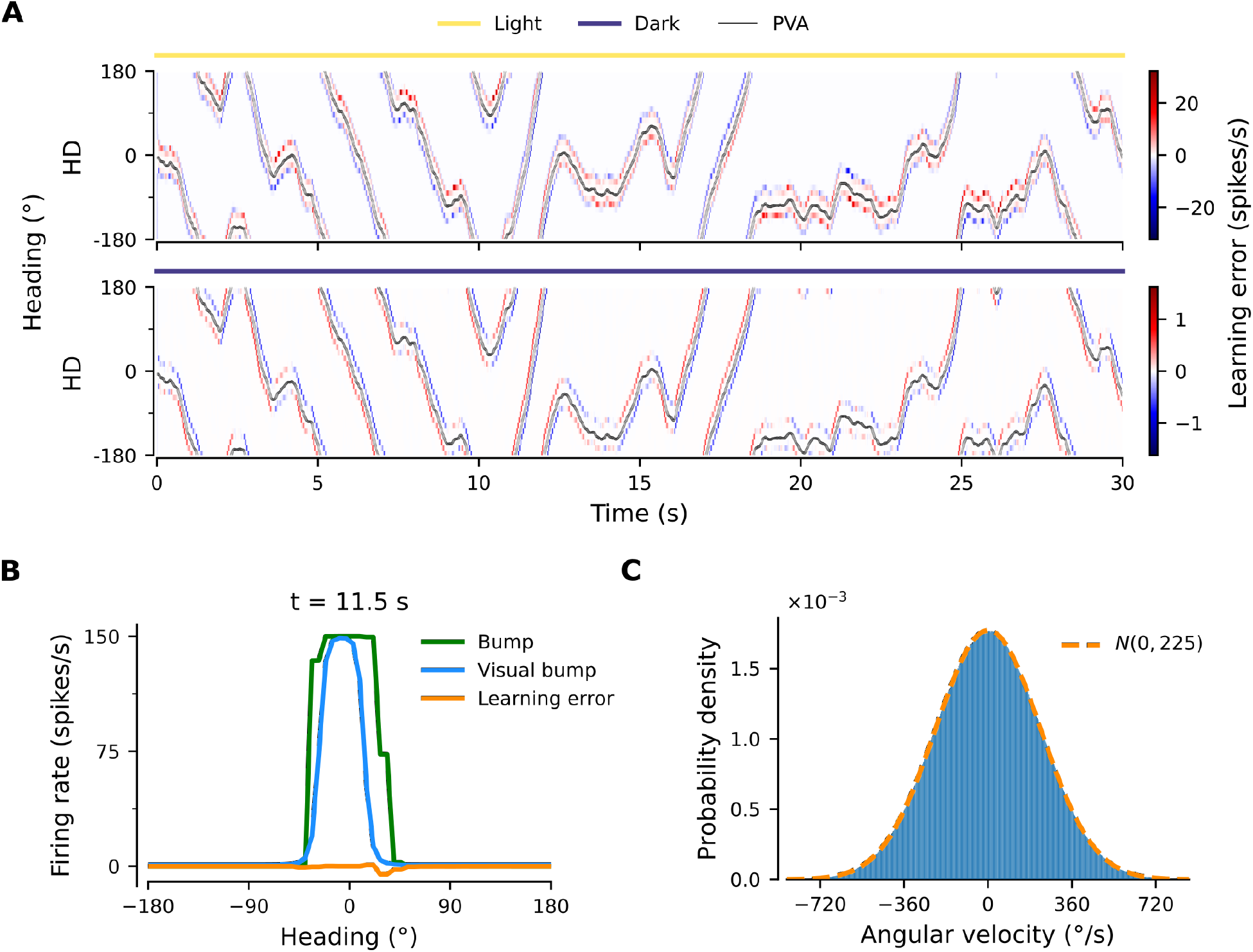
Details of learning. (A) Learning errors (eq. (17)) in converged network in light conditions (yellow overbars) or during PI in darkness (purple overbars). Note the different scales. (B) Snapshot of the bump, which has a square form, and the errors at t = 11.5 s in light conditions from (A). Also overlaid is the hypothetical form of the bump if only the visual input was present in the axon-proximal compartment of the HD neurons, termed “visual bump”. Notice that the errors are due to the fact that the visual bump is trailing in relation to the bump in the network. As a result, at the front of the bump the subthreshold visual input is actually inhibiting the bump. (C) Histogram of entrained velocities.

**Figure S3.**
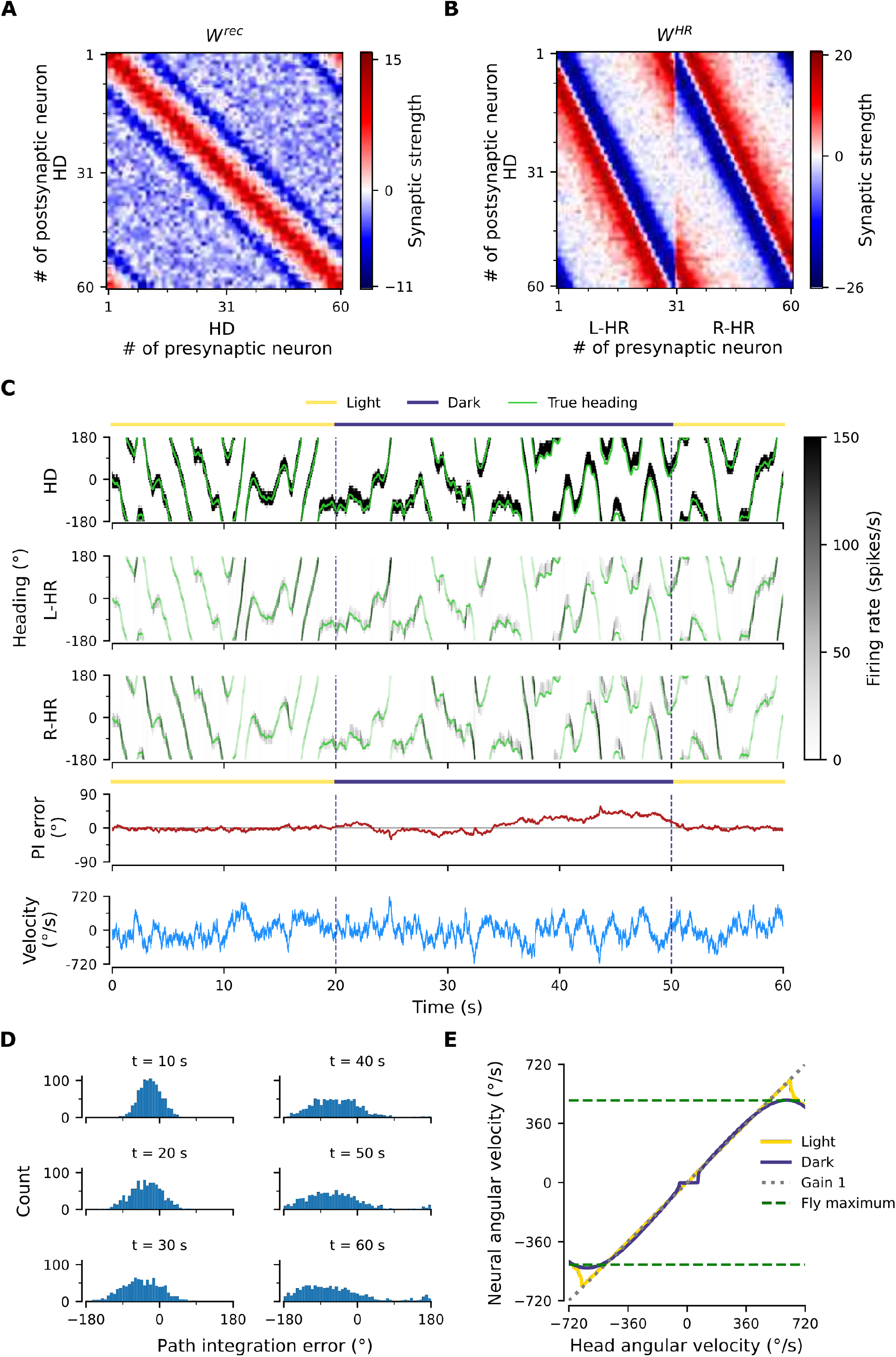
PI performance of perturbed network, after Gaussian noise with standard deviation ∼1.5 has been added to the synaptic connections in Fig. 3A,B. (A), (B) Resulting weight matrices after noise addition. (C) Example of PI. The activity of HD, L-HR and R-HR neurons along with the PI error and instantaneous angular velocity are displayed. (D) Temporal evolution of distribution of PI errors during PI in darkness. Compared to Fig. 2B the distribution gets wider faster, and also exhibits side bias. (E) PI is impaired compared to Fig. 2C, particularly for small angular velocities.

**Figure S4.**
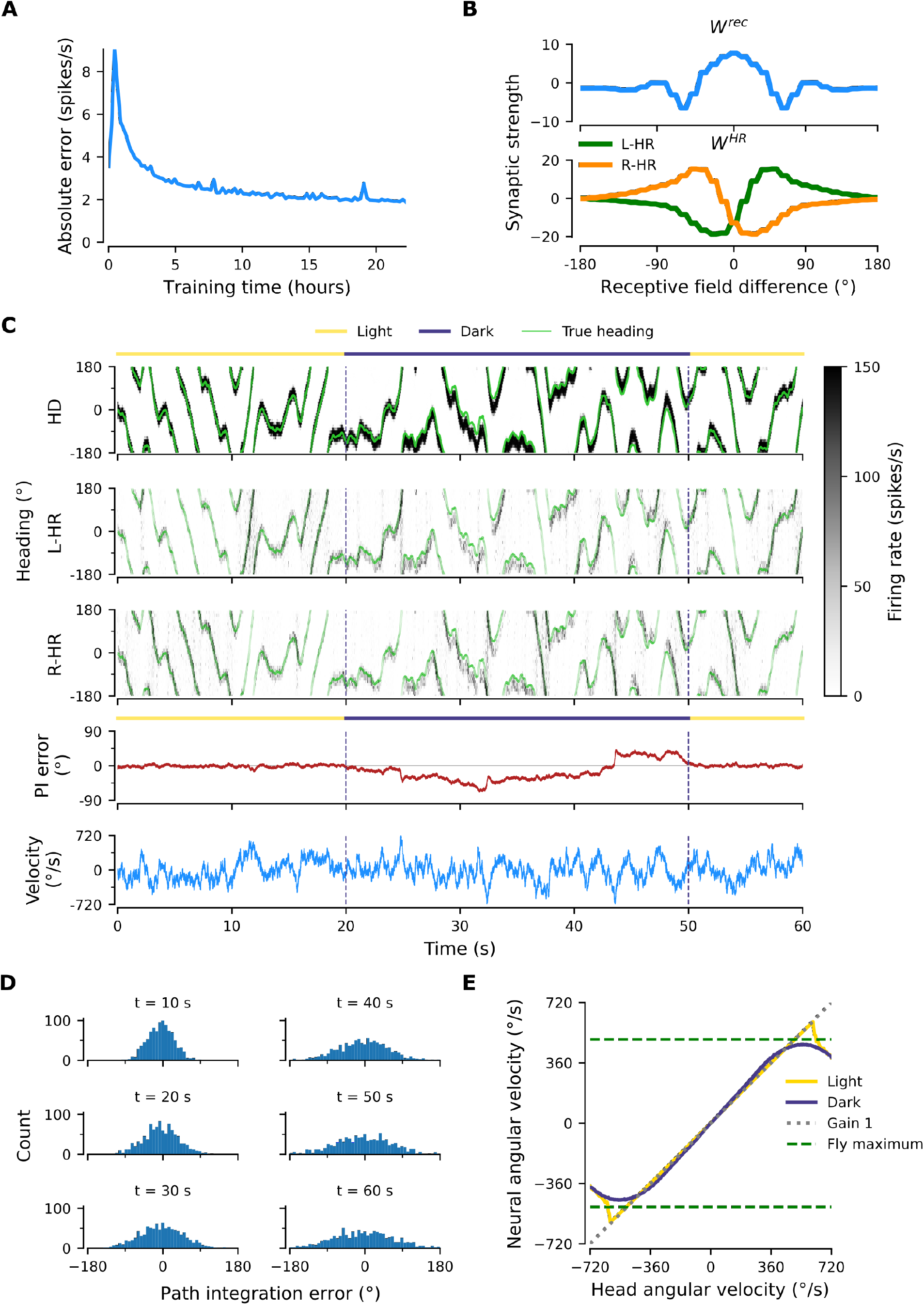
Learned network with *SNR=2*. (A) The mean learning error during training decreases and settles to a low value. (B) Profiles of learned weights. Both *W*^*rec*^ and *W*^*HR*^ (not shown) are circularly symmetric. Compared to Fig. 3C, the peak of the local excitatory profile in *W*^*rec*^ is not as pronounced, because the random noise corrupts the autocorrelations of HD neurons. (C) Example of PI. The activity of HD, L-HR and R-HR neurons along with the PI error and instantaneous angular velocity are displayed. The noise can be more readily seen in the HR activity, because HD cells not participating in the bump are deeply into inhibition. (D) Temporal evolution of distribution of PI errors during PI in darkness. Compared to Fig. 2B the distribution gets wider faster, however it does not exhibit side bias. (E) The network achieves almost perfect gain-1 PI, despite noisy inputs. Compared to Fig. 2C the performance is only slightly impaired.

**Figure S5.**
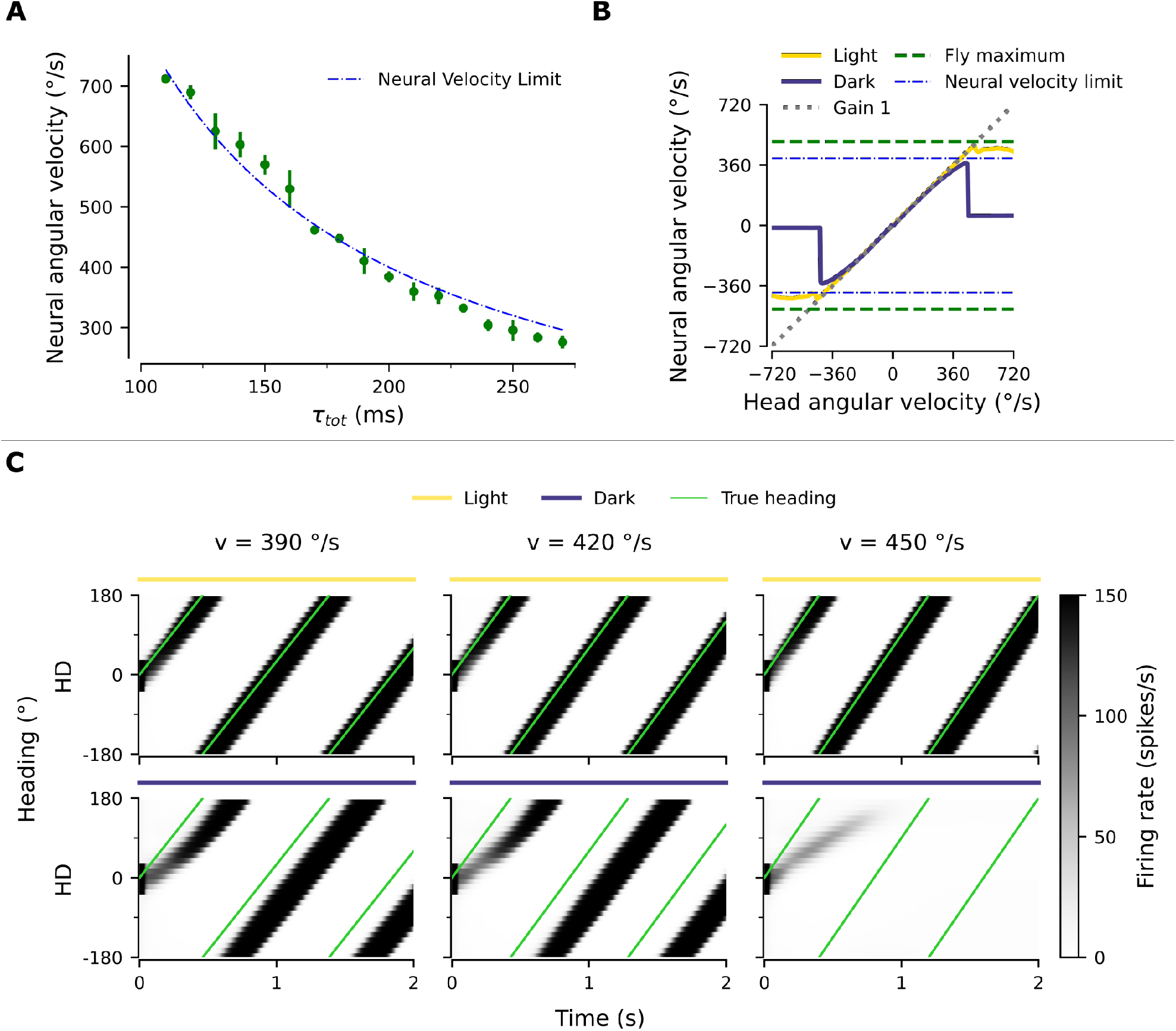
Network integrates angular velocities up to a limit set by synaptic delays. (A) Maximum neural velocity learned is inversely proportional to the total one-hop synaptic delays in the network. Green dots: point estimate of maximum neural velocity learned, green bars: 95 % confidence intervals (t-test). (B) Example neural velocity gain plot in network with increased synaptic delays (*τ*_*s*_ = 190 ms, *τ*_*l*_ = 10 ms). (C) Behavior of the network near the velocity limit. The example network in (B) is driven by a single velocity in every column, in light and darkness conditions. Near and below the limit, there is a delay in the appearance of the bump, which then path-integrates with near gain 1 in darkness. Above the limit however the bump cannot stabilize, resulting in the dip in neural velocity at the limit observed in (B).

## Mathematical Appendix

In this section we derive a reduced model for the dynamics of the synaptic weights during learning. The goal is to gain an intuitive understanding of the structure obtained in the full model (Figure 3 of the main text). Such a model reduction is obtained by 1) exploiting the circular symmetry in the system; 2) averaging weight changes across different speeds and moving directions; 3) writing dynamical equations in terms of convolutions and cross-correlations. With these methods, we derive a non-linear dynamical system for the weight changes as a function of head direction. Finally, we simulate this dynamical system and inspect how the different variables interact to obtain the final weights.

We study the learning equation (see Eqs. 11–15 in the main text where the low-pass filtering with time constant *τ*_*δ*_ has been ignored)

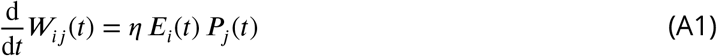

where

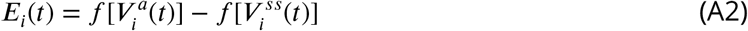

is the pre-synaptic error at cell *i* and

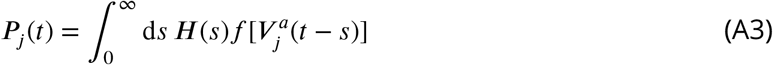

is the post-synaptic potential at HD cell, and *H* is a temporal filter (with time constants *τ*_*s*_ and *τ*_*l*_, see Eq. 13 of the main text).

### Clockwise movement

Assuming that the head turns clockwise (which equals to rightward rotation, i.e. rotation towards decreasing angles) and anti-clockwise (leftward, i.e. towards increasing angles) with equal probability, we can approximate the weight dynamics by summing the average weight change 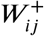 for clockwise movement and the average weight change 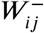 for anti-clockwise movement:

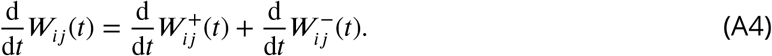

We start by assuming head movement at constant speed and we later generalize the results for multiple speeds. We compute the expected weight change 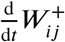 for one lap in the clockwise direction at speed *v*^+^ *>* 0:

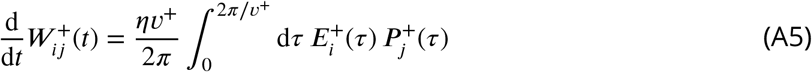

where

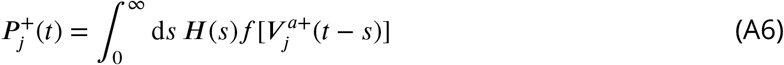

is the post-synaptic potential for clockwise movement, and

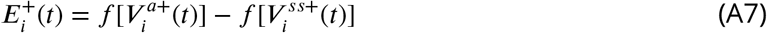

is the error for a clockwise movement. Assuming that the axon-proximal voltage is at steady state (Eq. 3 of the main text with the l.h.s. set to zero and 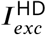 absorbed into *I*_*vis*_), the clockwise axon-proximal voltage reads

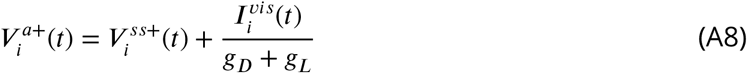

where (see Eq. 10 of the main text)

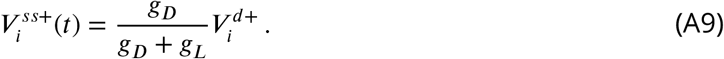

From Eqs. 1 and 2 of the main text, we can write the axon-distal voltage 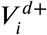 as a low-passfiltered version of the total axon-distal current 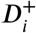 for clockwise movement (see also Eq. 13 of the main text):

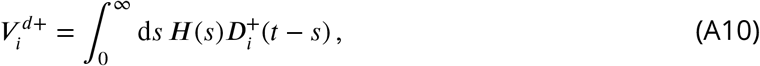

which yields

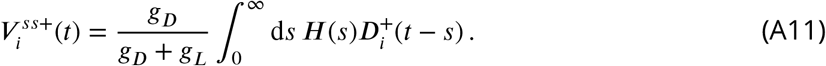

Importantly, the visual input *I*^*vis*^ is translation invariant:

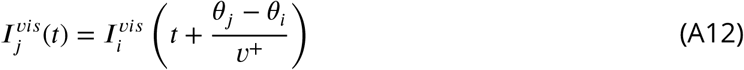

where *θ*_*j*_ and *θ*_*i*_ are the preferred head directions of cells *j* and *i*, respectively. As a result of this translation invariance, the recurrent weight matrix *W* develops circular symmetry:

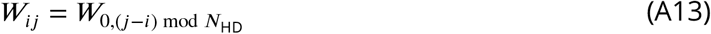

where *N*_HD_ is the number of HD cells in the system. Consequently, the post-synaptic potential 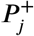 is also translation invariant:

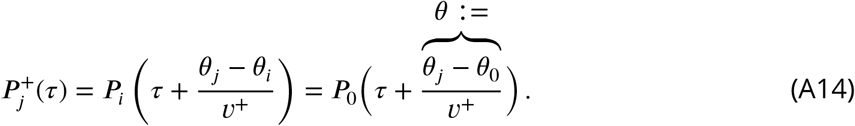

In this case, without loss of generality, we can rewrite Eq. A5 for a single row of the matrix 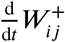 as a function of the angle difference *θ* = *θ*_*j*_ − *θ* _0_:

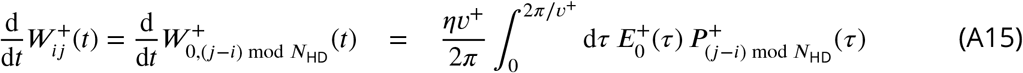

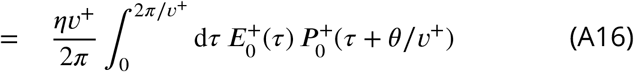

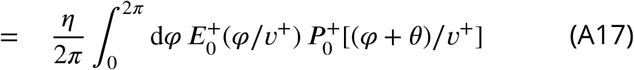

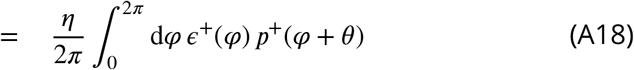

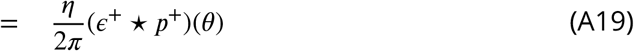

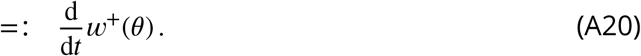

where we defined 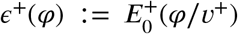 and 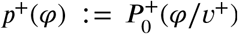, and ⋆ denotes circular cross-correlation.

From equation A6, we derive

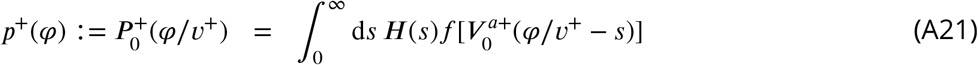

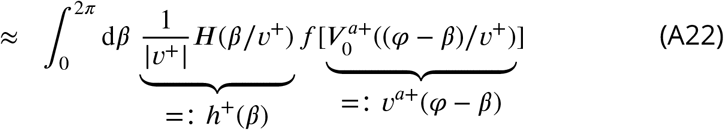

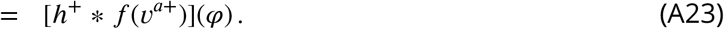

The approximation in Eq. A22 is valid if the temporal filter *H* is shorter than 2*π*/*v*^+^, that is for *H*(*t*) *«* 1 for *t >* 2*π*/*v*^+^, which holds for the filtering time constants and velocity distribution we assumed (Fig. A1). Therefore, plugging Eq. A23 into Eq. A20, we obtain:

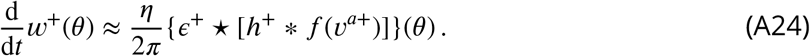

By using the definition of *ϵ*(φ)^+^ we derive

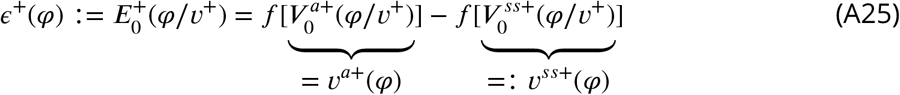

with (Eq. A8)

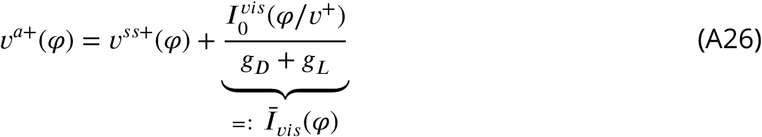

and (Eq. A11)

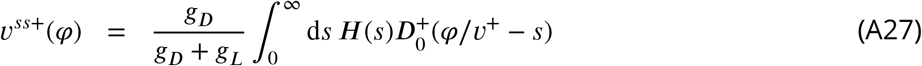

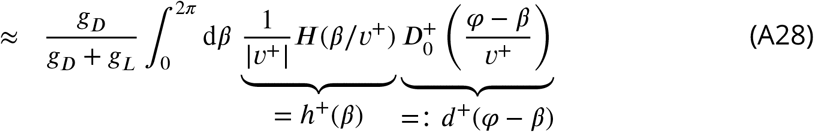

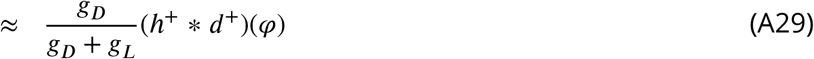

The approximation in Eq. A29 is valid if the temporal filter *H* is shorter than 2*π*/*v*^+^, which again holds true for our parameter choices (Fig. A1).

Calculation of the axon-distal input

Let us compute the axon-distal current 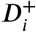 to neuron *i* for clockwise movement. From Eq. 1 of the main text, setting the l.h.s. to zero, and splitting the rotation-cell activities in the two populations (L-HR and R-HR), we derive

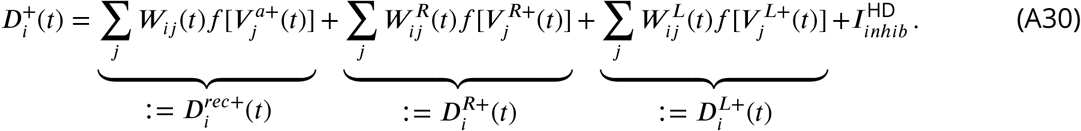

where 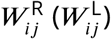 are the weights from the right (left) rotation cells, and 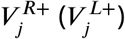 are the voltages of the right (left) rotation cells (see Eqs. 7–9 of the main text):

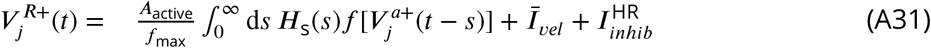

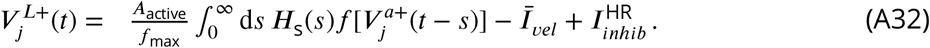

The function *H*_*S*_(*t*) = exp(−*t*/*τ*_*s*_)/*τ*_*s*_ is a temporal low pass filter with time constant *τ*_*s*_ and the velocity input reads (Eq. 9 of the main text)

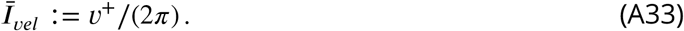

Eqs. A31 and A32, show that the rotation-cell voltages are re-scaled and filtered versions of the corresponding HD-cell firing rates with a baseline shift *Ī vel* that is differentially applied to right and left rotation cells.

From Eq. A30, we derive

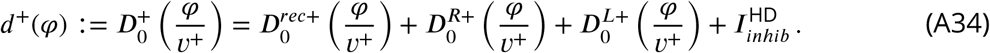

Assuming a large number _HD_ of HD cells evenly spaced around the circle, the recurrent axon-distal input reads

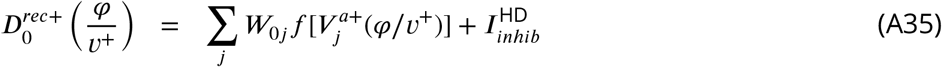

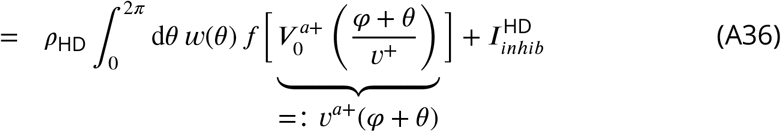

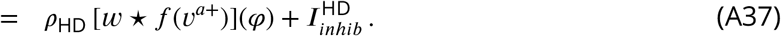

where *ρ*_HD_ = *N*_HD_/2*π* is the density of the HD neurons around the circle and we used the fact that the axon-proximal voltage is translation invariant (see also Eq A14):

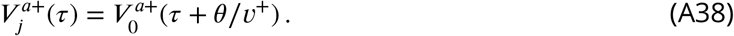

Following a similar procedure for 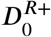 and 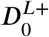, we obtain:

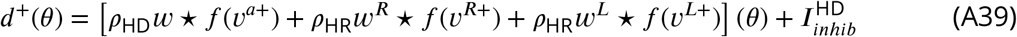

where *ρ*_HR_ = *N*_HR_/2*π* is the density of the HR neurons for one particular turning direction (note that we assumed *ρ*_HR_ = 2 *ρ*_HD_ in the main text). In deriving Eq. A39 we defined

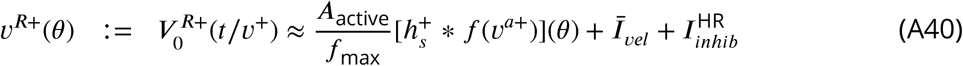

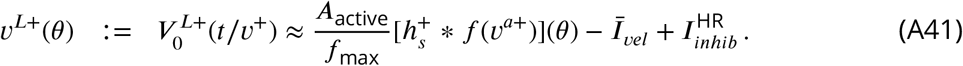

where we defined the filter 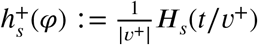, and the approximations are valid if *H*_*s*_(*t*/*v*^+^) *≪* 1 for *t >* 2 *π*/*v*^+^, which holds true for the time contant and velocity distribution assumed in the main text.

Finally, we compute the rotation-cells’ weights change. For these weights, the learning rule is the same as the one for the recurrent connections, except that the post-synaptic HD input is replaced by the post-synaptic HR input. Therefore, following the same procedure as in Eqs. A15-A23, the rotation weight changes are given by:

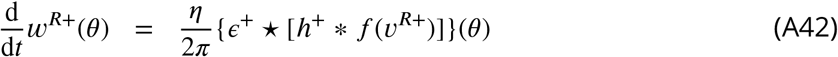

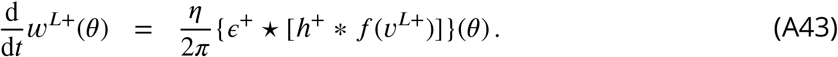

In summary, for clockwise movement, we obtain the following system of equations:

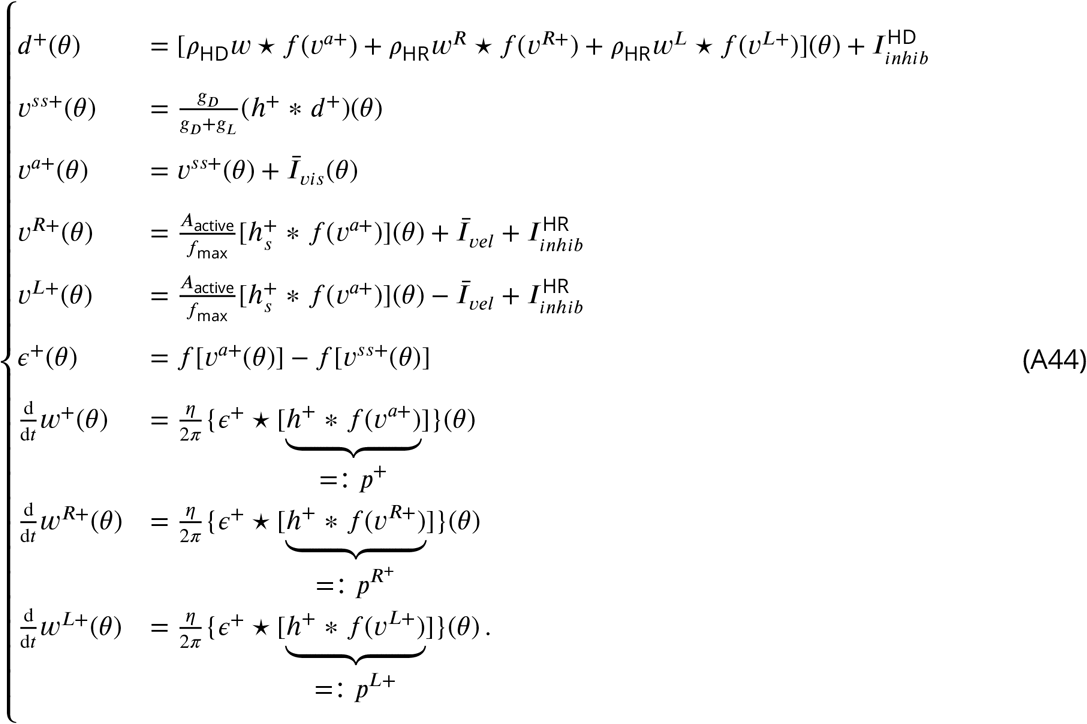

### Anti-clockwise movement

We now consider anticlockwise movements with speed *v*^-^ = −*v*^+^. First we note that the temporal filter

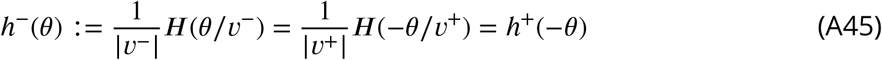

is a mirrored version about the origin of its clockwise counterpart *h*^+^, whereas the visual input is unchanged because it is symmetric around the origin (see Eq. 4 of the main text)

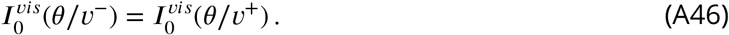

Let us first assume that

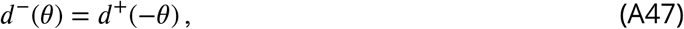

we shall verify the validity of this assumption self-consistently at the end of this section. From Eqs. A45–A47 it follows that 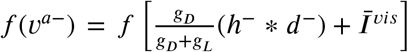 is a mirrored version of *f* (*v*^*a*+^), that is,

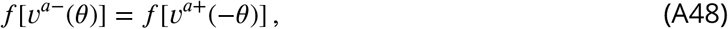

and, as a result,

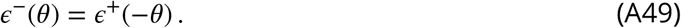

We now compute the anticlockwise weight change for the recurrent weights

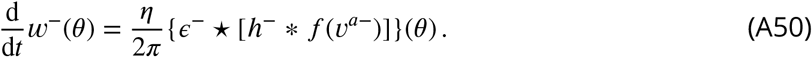

The r.h.s. of Eq. A50, without the *η*/(2*π*) pre-factor reads:

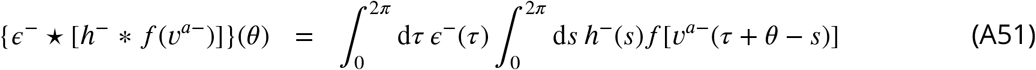

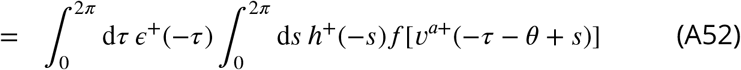

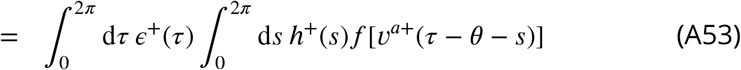

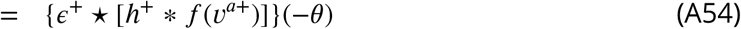

where from Eq. A52 to Eq. A53 we used variable substitution. Therefore, the weight change for clockwise movement is the mirrored version around the origin of the weight change for anticlockwise movement:

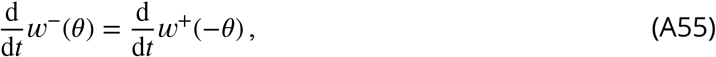

meaning that, with learning, the recurrent weights develop into an even function:

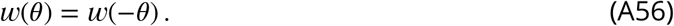

Let us now study the anticlockwise weight change for the rotation weights. The rotation-cell voltages during anticlockwise movement read:

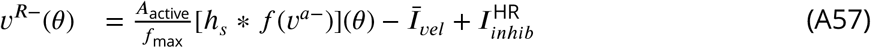

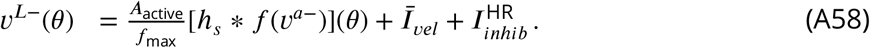

Using Eq. A48 in Eqs. A57 and A58 we find

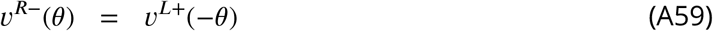

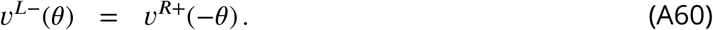

Therefore, applying the same procedure outlined in Eqs. A50–A54, to the anticlockwise change in the rotation weights yields

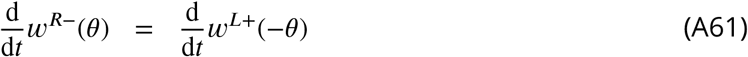

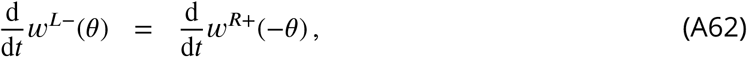

meaning that, during learning, the right and left rotation weights develop mirror symmetry:

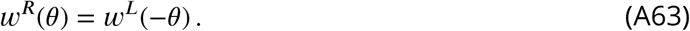

To verify that our original assumption in Eq. A47 holds, we compute the axon-distal input for anticlockwise movement:

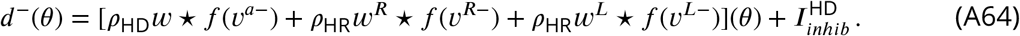

Using Eqs. A48, A56, A57, A58, A63 in Eq. A64, yields

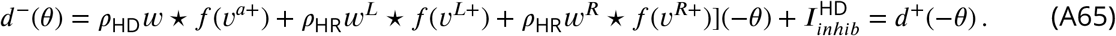

Finally, using Eqs. A55, A61, and A62, the total synaptic weight changes for both clockwise and anticlockwise movement read

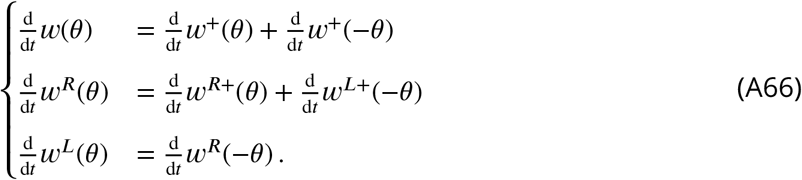

### Averaging across speeds

So far, we have only considered head turnings at a fixed speed *v*^+^ (clockwise) and *v*^-^ = −*v*^+^ (anticlockwise). However, in the full model described in the main text, velocities are sampled stochastically from an OU process. This random process generates a half-normal distribution of speeds with spread *σ*_*v*_/2 (Fig. A1, left, see also Table 1 in the main text). We thus compute the expected weight changes with respect to this speed distribution:

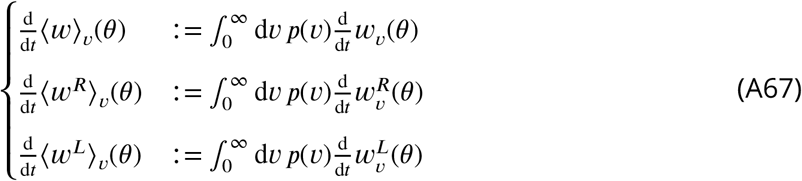

where *w*_*v*_ is the weight change for speed |*v*^+^| = |*v*^-^ |= *v* and *p*(*v*) is an half-normal distribution with spread *σ*_*v*_/2.

### Simulation of the reduced model

In this section, we show the dynamics of the reduced model numerically simulated according to Eqs. A44, A66, and A67. Weight changes are computed at discrete time steps and integrated using the forward Euler method. At each time step we compute the weight changes for each speed *v* (Eqs. A44 and A66) and we estimate the expected weight change according to Eq. A67. We then update the weights and proceed to the next step of the simulation. Note that Eq. A44 requires the firing rates of HD and HR cells at the previous time step (recurrent input, first line of Eq. A44). Therefore, at each time step, we save the HD and HR firing rates for every speed value *v* and provide them as input to the next iteration of the simulation.

**Figure A1.**
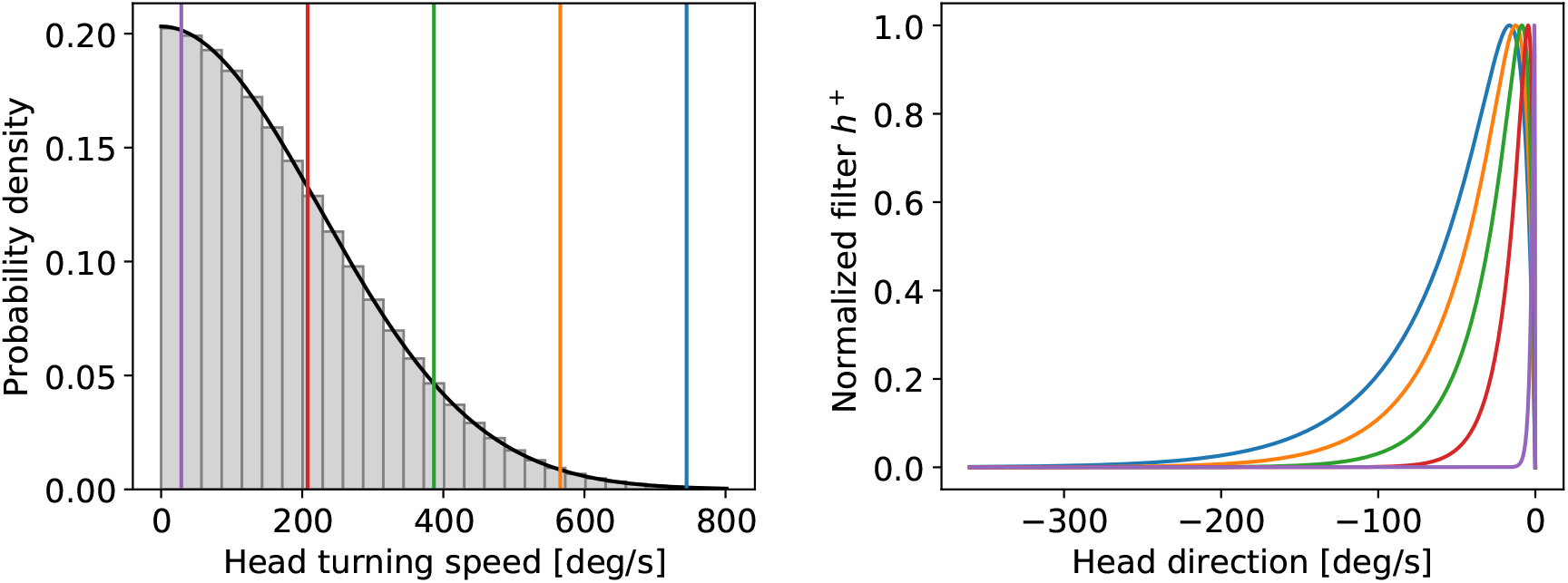
Left: assumed distribution of head-turning speeds (black) and discrete approximation used for the simulations. The colored vertical lines indicate speeds for which the filter *h*^+^ is plotted in the right panel. Right: temporal filter *h*^+^(*θ*) for several example speeds (see vertical lines in the left panel). Note that even for the largest speeds (blue curve) the filter decays within one turn around the circle.

Fig. A2 shows the evolution of the reduced system for 400 time steps, starting from an initial condition where all weights are zero. One can see that from time steps 75 to 100 the system switches from a linear regime (HD firing rates below saturation, see top panel) to a non-linear regime (saturated HD rates). Such a switch is accompanied by peaks in the average absolute error (third panel from the top). Notably, the rotation weights start developing a structure only after such switch has occurred (see two bottom panels)—a feature that has been observed also in the full model (Fig. 3e of the main text).

#### Development of the recurrent weights

Figure A3 provides an intuitive explanation for the shape of the recurrent-weight profiles *w* that emerge during learning. The first column shows the evolution of the recurrent weights in the linear regime (*t* = 25), i.e., before the HD rates reach saturation. In this regime, both recurrent and rotation weights are small, and the steady-state axon-distal rate

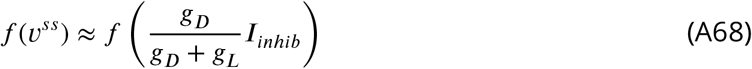

is flat and close to zero. Therefore, the HD output rate *f* (*v*^*a*^) is dominated by the visual input *Ī* _*vis*_ (Eq. A44, third line), which has the shape of a localized bump (panel A1). Thus the error *ϵ* has also the shape of a bump (B1). Additionally, the post-synaptic inputs *p*^+^ and *p*^−^ are shifted and filtered versions of this bump (Eq. A44, seventh line). The recurrent weight changes d*w*^+^ and d*w*^−^ for clockwise and anticlockwise movement are given by the cross-correlation of the errors *ϵ*^+^ and *ϵ*^−^ with the post-synaptic inputs *p*^+^ and *p*^−^ (panel C1; see Eq. A44 seventh line and Eq. A50). Note that because *a*(*x*) ⋆ *b*(*x*) = *a*(− *x*) * *b*(*x*), the operation of cross-correlation can be understood graphically as a convolution between the mirrored first function *a* and the second function *b*. Such a mirroring is irrelevant in C1 (linear regime) because the error is an even function, but becomes important in C2 (non-linear regime). As a result of this cross-correlation, the recurrent recurrent-weight changes d*w*^+^ and d*w*^−^ are shifted bumps (colored lines in C1), which merge into a single central bump after summing clockwise and anticlockwise contributions (black line in C1). Therefore, in the linear regime, the recurrent weights develop a single central peak in the origin (panel D1).

**Figure A2.**
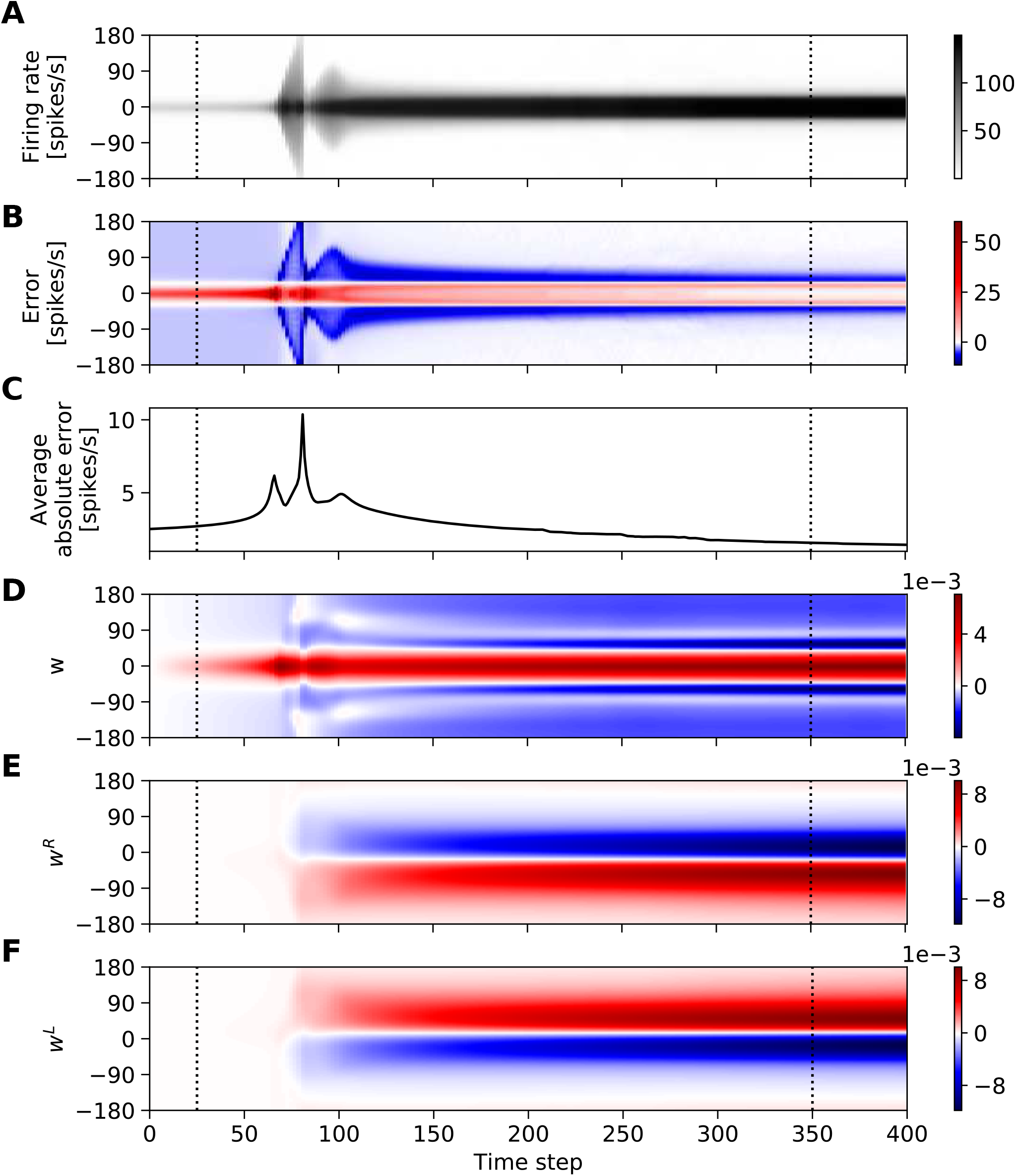
Evolution of the reduced model. The figure shows from top to bottom: A) the HD-cells’ firing rate *f* (*v*^*a*+^); B) the error *ϵ*; C) the average absolute error; D) the recurrent weights *w*; E-F) the rotation weights *w*^*R*^ and *w*^*L*^. The HD firing rate and the errors (panels A-C) are averaged across speeds and and both movement directions. The vertical dashed lines denote the time points shown in Figs. A3 and A4.

The second column of Fig. A3 shows the development of the recurrent weights in the non-linear regime (time step 350). Panel A2 shows that in this scenario the HD firing-rate bumps are broader and approach saturation due to the strong recurrent input. The coupling between the axon-distal and axon-proximal compartment acts as a self-amplifying signal during learning which results in the activity of all active neurons participating in the bump reaching saturation. Additionally, because the recurrent input is filtered in time (Eq. A44, second line), such bumps are also shifted towards the direction of movement. Importantly, due to the lack of visual input, within the receptive field the steady-state axon-distal rates are always smaller than the firing rates. As a result, the errors *ϵ*^+^ and *ϵ*^−^ show small negative bumps in the direction of movement, and small positive bumps in the opposite direction (panel B2). Additionally, the post-synaptic inputs *p*^+^ and *p*^−^ shift further apart from the origin. Consequently, the total weight change d*w* develops negative peaks around 60 degrees (black line in C2, contrast to panel C1), and these peaks get imprinted in the final recurrent weights’ profiles (panel D2).

#### Development of the rotation weights

Figure A4 provides an intuitive explanation for the shape of the rotation-weights profiles *w*^*R*^ and *w*^*L*^ that emerge during learning. The first column shows the evolution of the rotation weights in the linear regime (*t* = 25), i.e., before the HD rates reach saturation. In this regime, the rotation-cell firing rates are filtered versions of the HD bumps but re-scaled by a factor *A*_active_/*f*_max_ ≈ 0.013 and baseline-shifted by an amount 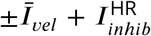 (Eq. A44 lines 4 and 5; panel A1, compare to Fig. A3A1. Panel B1 shows that the errors *ϵ*^+^ and *ϵ*^−^ overlap and have the shape of a bump centered at the origin (same curves as in Figure A3B1). Additionally, the post-synaptic potentials *p*^*R*±^ and *p*^*L*±^ in B1 are filtered versions of the curves in A1 (Eq. A44, lines 7 and 8). As a result, the weight changes d*w* ^*R*±^ and d*w*^*L*±^, i.e., the errors cross-correlated by the post-synaptic potentials, appear similar to the bumps in A1, but they are smoother and further apart from the origin (panel C1). Finally, such weight changes get imprinted in the rotation weights (panel D1).

The second column shows the evolution of the rotation weights in the non-linear regime (*t* = 350), i.e., after the HD rates reach saturation. In this case, the large recurrent input gives rise to larger rotation rates (A2, compare to A1) and larger post-synaptic potentials (B2, com-pare to B1). In panel B2, we can see that the errors *ϵ*^+^ and *ϵ*^−^ show positive and negatives peaks shifted from the origin (same curves as in Figure A3B2), which generate weight changes with both positive and negative lobes (panel C2). Such weight changes get finally imprinted in the rotation weights (panel D2).

**Figure A3.**
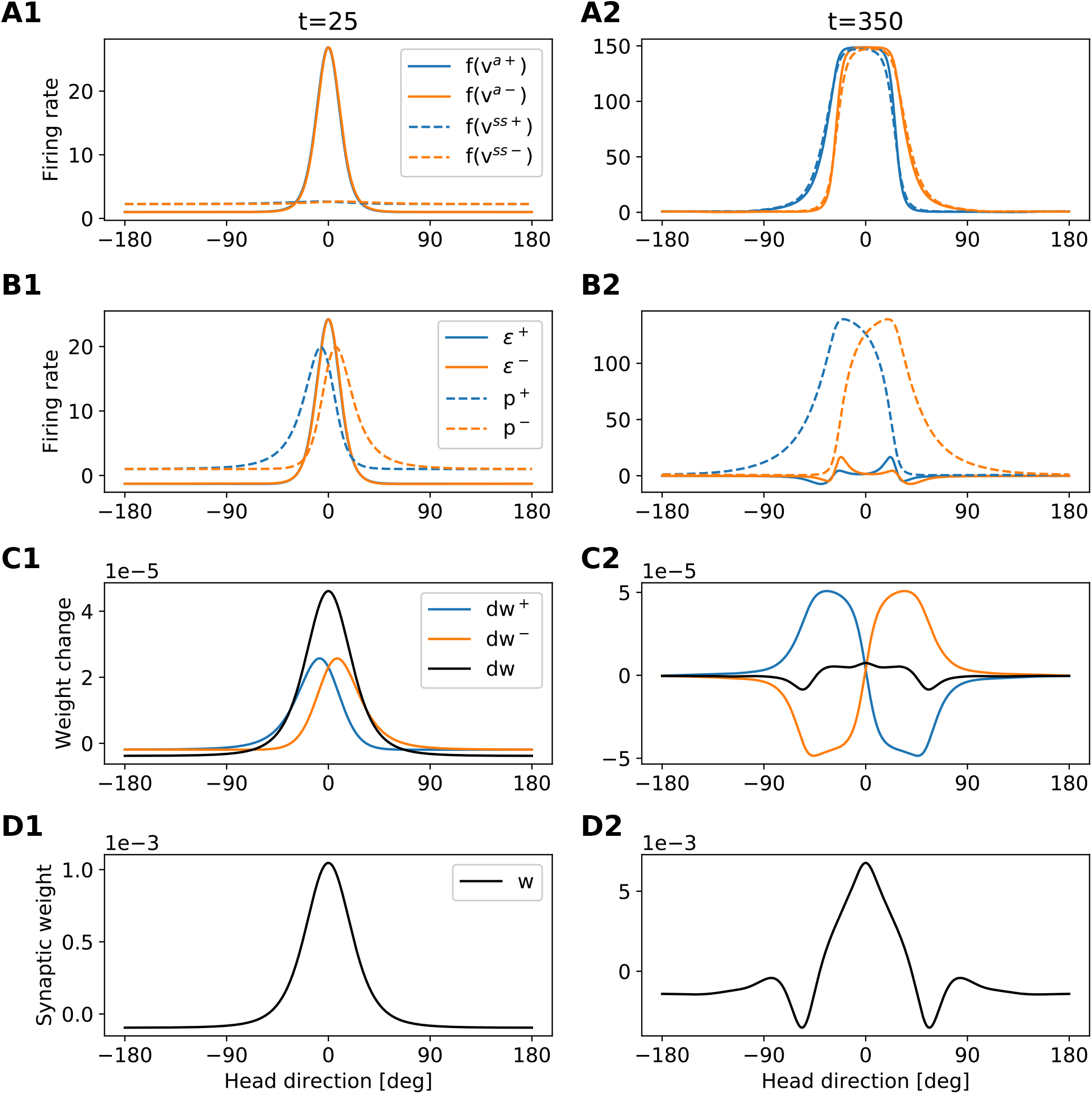
Development of the recurrent weights. The figure provides an intuition for the shape of the recurrent-weights profiles that emerge during learning. Each column refers to a different time step (see also dashed lines in Fig A2). Each row shows a different set of variables of the model (see legends in the first column). The figure is to be read from top to bottom, because variables in the lower rows are computed from variables in the upper rows. Blue (orange) lines always refer to clockwise (anticlockwise) motion. Black lines in C show the total weight changes for both clockwise and anti-clockwise motion, i.e., d*w* = d*w*^+^ + d*w*^−^.

**Figure A4.**
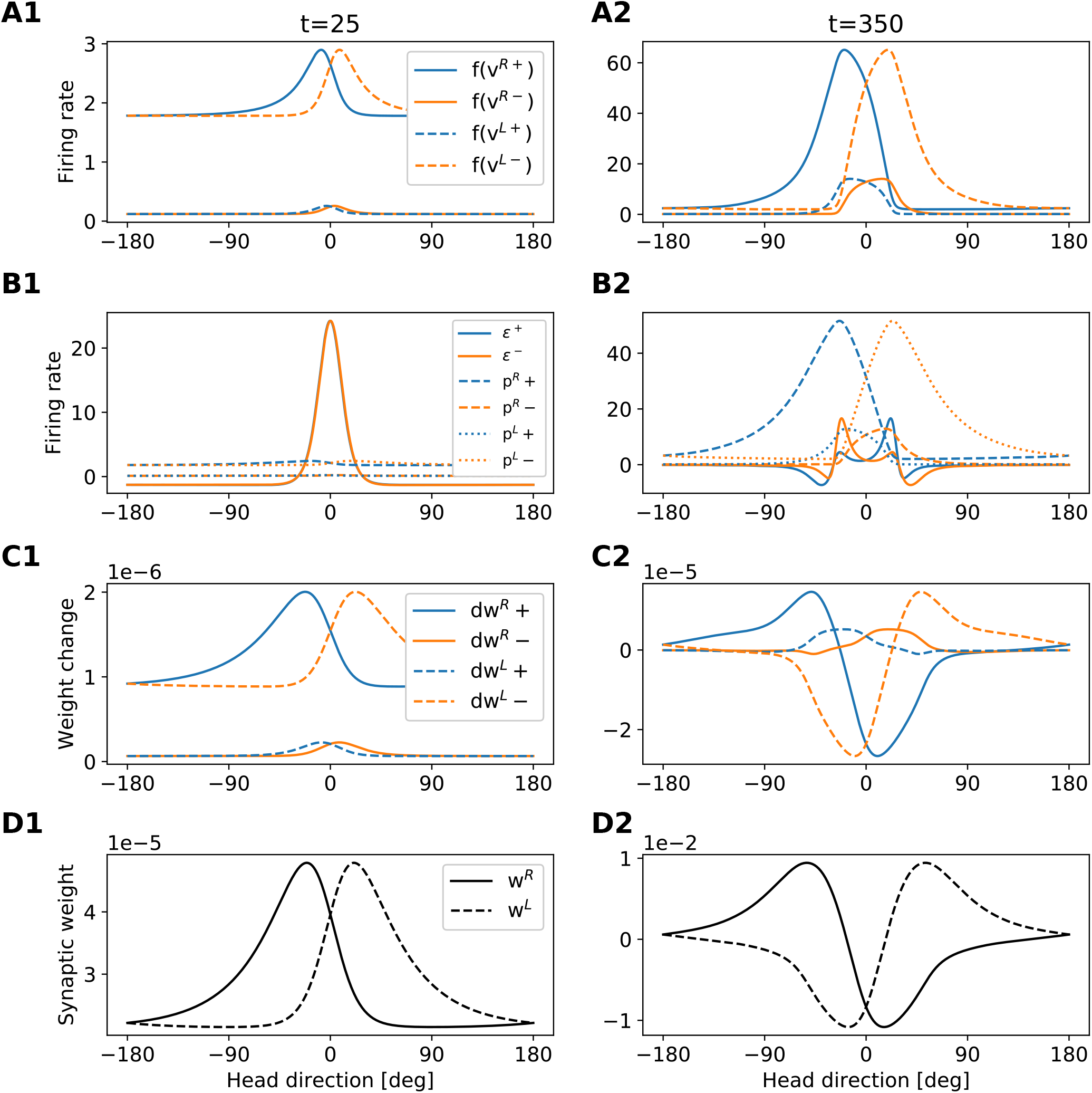
Development of the rotation weights. The figure provides an intuition for the shape of the rotation-weights profiles that emerge during learning. Each column refers to a different time step (see also dashed lines in Fig A2). Each row shows a different set of variables of the model (see legends in the first column). The figure is to be read from top to bottom, because variables in the lower rows are computed from variables in the upper rows. Blue (orange) lines always refer to clockwise (anticlockwise) motion.

## References

Abbott, L. F., Bock, D. D., Callaway, E. M., Denk, W., Dulac, C., Fairhall, A. L., Fiete, I., Harris, K. M., Helm-staedter, M., Jain, V., Kasthuri, N., LeCun, Y., Lichtman, J. W., Littlewood, P. B., Luo, L., Maunsell, J. H., Reid, R. C., Rosen, B. R., Rubin, G. M., Sejnowski, T. J., Seung, H. S., Svoboda, K., Tank, D. W., Tsao, D., and Van Essen, D. C. (2020). The mind of a mouse. Cell, 182(6):1372–1376.

Amari, S. (1977). Dynamics of pattern formation in lateral-inhibition type neural fields. Biological Cybernetics, 27(2):77–87.

Bell, C. C., Han, V. Z., Sugawara, Y., and Grant, K. (1997). Synaptic plasticity in a cerebellum-like structure depends on temporal order. Nature, 387(6630):278–281.

Ben-Yishai, R., Bar-Or, R. L., and Sompolinsky, H. (1995). Theory of orientation tuning in visual cortex. Proceedings of the National Academy of Sciences of the United States of America, 92(9):3844–3848.

Beniaguev, D., Segev, I., and London, M. (2019). Single cortical neurons as deep artificial neural networks. bioRxiv.

Brea, J., Gaál, A. T., Urbanczik, R., and Senn, W. (2016). Prospective coding by spiking neurons. PLOS Computational Biology, 12(6):e1005003.

Burak, Y. and Fiete, I. R. (2009). Accurate path integration in continuous attractor network models of grid cells. PLoS Computational Biology, 5(2):e1000291.

Burak, Y. and Fiete, I. R. (2012). Fundamental limits on persistent activity in networks of noisy neurons. Proceedings of the National Academy of Sciences of the United States of America, 109(43):17645–17650.

Carpenter, G. A. and Grossberg, S. (1987). A massively parallel architecture for a self-organizing neural pattern recognition machine. Computer Vision, Graphics, and Image Processing, 37(1):54–115.

Chaudhuri, R., Gerçek, B., Pandey, B., Peyrache, A., and Fiete, I. (2019). The intrinsic attractor manifold and population dynamics of a canonical cognitive circuit across waking and sleep. Nature Neuroscience, 22(9):1512–1520.

Clements, J., Dolafi, T., Umayam, L., Neubarth, N. L., Berg, S., Scheffer, L. K., and Plaza, S. M. (2020). neuPrint: Analysis tools for EM connectomics. bioRxiv.

D’Albis, T. and Kempter, R. (2020). Recurrent amplification of grid-cell activity. Hippocampus, 30(12):1268–1297.

Darwin, C. (1873). Origin of certain instincts. Nature, 7(179):417–418.

DePasquale, B., Cueva, C. J., Rajan, K., Escola, G. S., and Abbott, L. F. (2018). full-FORCE: A target-based method for training recurrent networks. PLOS ONE, 13(2):e0191527.

Doron, G., Shin, J. N., Takahashi, N., Drüke, M., Bocklisch, C., Skenderi, S., de Mont, L., Toumazou, M., Ledderose, J., Brecht, M., Naud, R., and Larkum, M. E. (2020). Perirhinal input to neocortical layer 1 controls learning. Science, 370(6523).

Eichenbaum, H. (2017). The role of the hippocampus in navigation is memory. Journal of Neurophysiology, 117(4):1785–1796.

Etienne, A. S., Maurer, R., and Séguinot, V. (1996). Path integration in mammals and its interaction with visual landmarks. Journal of experimental biology, 199(1):201–209.

Fisher, Y. E., Lu, J., D’Alessandro, I., and Wilson, R. I. (2019). Sensorimotor experience remaps visual input to a heading-direction network. Nature, 576(7785):121–125.

Franconville, R., Beron, C., and Jayaraman, V. (2018). Building a functional connectome of the drosophila central complex. Elife, 7:e37017.

Gallistel, C. R. (1993). The Organization of Learning. Bradford Books/MIT Press.

Geurten, B. R. H., Jähde, P., Corthals, K., and Göpfert, M. C. (2014). Saccadic body turns in walking drosophila. Frontiers in Behavioral Neuroscience, 8.

Gidon, A., Zolnik, T. A., Fidzinski, P., Bolduan, F., Papoutsi, A., Poirazi, P., Holtkamp, M., Vida, I., and Larkum, M. E. (2020). Dendritic action potentials and computation in human layer 2/3 cortical neurons. Science, 367(6473):83–87.

Goldman, M., Compte, A., and Wang, X.-J. (2009). Neural integrator models.In Encyclopedia of Neuroscience, pages 165–178. Elsevier.

Gouwens, N. W. and Wilson, R. I. (2009). Signal propagation in drosophila central neurons. Journal of Neuroscience, 29(19):6239–6249.

Green, J., Adachi, A., Shah, K. K., Hirokawa, J. D., Magani, P. S., and Maimon, G. (2017). A neural circuit architecture for angular integration in drosophila. Nature, 546(7656):101–106.

Green, J., Vijayan, V., Pires, P. M., Adachi, A., and Maimon, G. (2019). A neural heading estimate is compared with an internal goal to guide oriented navigation. Nature Neuroscience, 22(9):1460–1468.

Guerguiev, J., Lillicrap, T. P., and Richards, B. A. (2017). Towards deep learning with segregated dendrites. Elife, 6.

Hafting, T., Fyhn, M., Molden, S., Moser, M.-B., and Moser, E. I. (2005). Microstructure of a spatial map in the entorhinal cortex. Nature, 436(7052):80–806.

Hahnloser, R. H. R. (2003). Emergence of neural integration in the head-direction system by visual supervision. Neuroscience, 120(3):877–891.

Itti, L., Koch, C., and Niebur, E. (1998). A model of saliency-based visual attention for rapid scene analysis. IEEE Transactions on Pattern Analysis and Machine Intelligence, 20(11):1254–1259.

Jayakumar, R. P., Madhav, M. S., Savelli, F., Blair, H. T., Cowan, N. J., and Knierim, J. J. (2019). Recalibration of path integration in hippocampal place cells. Nature, 566(7745):533–537.

Kilpatrick, Z. P., Ermentrout, B., and Doiron, B. (2013). Optimizing working memory with heterogeneity of recurrent cortical excitation. Journal of Neuroscience, 33(48):18999–19011.

Kim, S. S., Hermundstad, A. M., Romani, S., Abbott, L. F., and Jayaraman, V. (2019). Generation of stable heading representations in diverse visual scenes. Nature, 576(7785):126–131.

Kim, S. S., Rouault, H., Druckmann, S., and Jayaraman, V. (2017). Ring attractor dynamics in theDrosophilacentral brain. Science, 356(6340):849–853.

Lake, B. M., Ullman, T. D., Tenenbaum, J. B., and Gershman, S. J. (2016). Building machines that learn and think like people. Behavioral and Brain Sciences, 40.

Langston, R. F., Ainge, J. A., Couey, J. J., Canto, C. B., Bjerknes, T. L., Witter, M. P., Moser, E. I., and Moser, M.-B. (2010). Development of the spatial representation system in the rat. Science, 328(5985):1576– 1580.

Larkum, M. (2013). A cellular mechanism for cortical associations: an organizing principle for the cerebral cortex. Trends in Neurosciences, 36(3):141–151.

Larkum, M. E., Zhu, J. J., and Sakmann, B. (1999). A new cellular mechanism for coupling inputs arriving at different cortical layers. Nature, 398(6725):338–341.

McNaughton, B. L., Barnes, C. A., Gerrard, J. L., Gothard, K., Jung, M. W., Knierim, J. J., Kudrimoti, H., Qin, Y., Skaggs, W., Suster, M., et al. (1996). Deciphering the hippocampal polyglot: the hippocampus as a path integration system. Journal of Experimental Biology, 199(1):173–185.

Mittelstaedt, M. L. and Mittelstaedt, H. (1980). Homing by path integration in a mammal. Naturwissenschaften, 67(11):566–567.

Mizumori, S. and Williams, J. (1993). Directionally selective mnemonic properties of neurons in the lateral dorsal nucleus of the thalamus of rats. Journal of Neuroscience, 13(9):4015–4028.

Moser, E. I., Kropff, E., and Moser, M.-B. (2008). Place cells, grid cells, and the brain’s spatial representation system. Annual Review of Neuroscience, 31(1):69–89.

O’Keefe, J., Nadel, L., and of Psychology Lynn Nadel, R.P. (1978). The Hippocampus as a Cognitive Map. Oxford University Press.

Omoto, J. J., Keleş, M. F., Nguyen, B.-C. M., Bolanos, C., Lovick, J. K., Frye, M. A., and Hartenstein, V. (2017). Visual input to the drosophila central complex by developmentally and functionally distinct neuronal populations. Current Biology, 27(8):1098–1110.

Page, H. J. I., Walters, D., and Stringer, S. M. (2018). A speed-accurate self-sustaining head direction cell path integration model without recurrent excitation. Network: Computation in Neural Systems, 29(1-4):37–69.

Poirazi, P., Brannon, T., and Mel, B. W. (2003). Pyramidal neuron as two-layer neural network. Neuron, 37(6):989–999.

Quirk, G. J., Muller, R. U., and Kubie, J. L. (1990). The firing of hippocampal place cells in the dark depends on the rat’s recent experience. Journal of neuroscience, 10(6):2008–2017.

Raccuglia, D., Huang, S., Ender, A., Heim, M.-M., Laber, D., Suárez-Grimalt, R., Liotta, A., Sigrist, S. J., Geiger, J. R., and Owald, D. (2019). Network-specific synchronization of electrical slow-wave oscillations regulates sleep drive in drosophila. Current Biology, 29(21):3611–3621.e3.

Ranck, J. B. (1984). Head direction cells in the deep layer of dorsal presubiculum in freely moving rats. Society of Neuroscience Abstract, 10:599.

Redish, A. D., Elga, A. N., and Touretzky, D. S. (1996). A coupled attractor model of the rodent head direction system. Network: Computation in Neural Systems, 7(4):671–685.

Samsonovich, A. and McNaughton, B. L. (1997). Path integration and cognitive mapping in a continuous attractor neural network model. Journal of Neuroscience, 17(15):5900–5920.

Seelig, J. D. and Jayaraman, V. (2015). Neural dynamics for landmark orientation and angular path integration. Nature, 521(7551):186–191.

Seung, H. S. (1996). How the brain keeps the eyes still. Proceedings of the National Academy of Sciences of the United States of America, 93(23):13339–13344.

Skaggs, W. E., Knierim, J. J., Kudrimoti, H. S., and McNaughton, B. L. (1995). A model of the neural basis of the rat’s sense of direction. Advances in neural information processing systems, 7:173–180.

Song, P. and Wang, X.-J. (2005). Angular path integration by moving “hill of activity”: A spiking neuron model without recurrent excitation of the head-direction system. Journal of Neuroscience, 25(4):1002– 1014.

Stowers, J. R., Hofbauer, M., Bastien, R., Griessner, J., Higgins, P., Farooqui, S., Fischer, R. M., Nowikovsky, K., Haubensak, W., Couzin, I. D., Tessmar-Raible, K., and Straw, A. D. (2017). Virtual reality for freely moving animals. Nature Methods, 14(10):995–1002.

Stringer, S. M., Trappenberg, T. P., Rolls, E. T., and de Araujo, I. E. T. (2002). Self-organizing continuous attractor networks and path integration: one-dimensional models of head direction cells. Network: Computation In Neural Systems, 13(2):217–242.

Tolman, E. C. (1948). Cognitive maps in rats and men. Psychological Review, 55(4):189–208.

Turner-Evans, D., Wegener, S., Rouault, H., Franconville, R., Wolff, T., Seelig, J. D., Druckmann, S., and Jayaraman, V. (2017). Angular velocity integration in a fly heading circuit. eLife, 6:e23496.

Turner-Evans, D. B., Jensen, K. T., Ali, S., Paterson, T., Sheridan, A., Ray, R. P., Wolff, T., Lauritzen, J. S., Rubin, G. M., Bock, D. D., and Jayaraman, V. (2020). The neuroanatomical ultrastructure and function of a biological ring attractor. Neuron, 108(1):145–163.e10.

Tuthill, J. C. (2009). Lessons from a compartmental model of a drosophila neuron. Journal of Neuroscience, 29(39):12033–12034.

Urbanczik, R. and Senn, W. (2014). Learning by the dendritic prediction of somatic spiking. Neuron, 81(3):521–528.

Wang, X.-J. (2001). Synaptic reverberation underlying mnemonic persistent activity. Trends in Neurosciences, 24(8):455–463.

Wang, X.-J. (2002). Probabilistic decision making by slow reverberation in cortical circuits. Neuron, 36(5):955–968.

Xie, X., Hahnloser, R. H. R., and Seung, H. S. (2002). Double-ring network model of the head-direction system. Physical Review E, 66(4):041902.

Xie, X. and Seung, H. S. (2000). Spike-based learning rules and stabilization of persistent neural activity. Advances in neural information processing systems, 12:199–208.

Xu, C. S., Januszewski, M., Lu, Z., ya Takemura, S., Hayworth, K. J., Huang, G., Shinomiya, K., MaitinShepard, J., Ackerman, D., Berg, S., Blakely, T., Bogovic, J., Clements, J., Dolafi, T., Hubbard, P., Kain-mueller, D., Katz, W., Kawase, T., Khairy, K. A., Leavitt, L., Li, P. H., Lindsey, L., Neubarth, N., Olbris, D. J., Otsuna, H., Troutman, E. T., Umayam, L., Zhao, T., Ito, M., Goldammer, J., Wolff, T., Svirskas, R., Schlegel, P., Neace, E. R., Knecht, C. J., Alvarado, C. X., Bailey, D. A., Ballinger, S., Borycz, J. A., Canino, B. S., Cheatham, N., Cook, M., Dreher, M., Duclos, O., Eubanks, B., Fairbanks, K., Finley, S., Forknall, N., Francis, A., Hopkins, G. P., Joyce, E. M., Kim, S., Kirk, N. A., Kovalyak, J., Lauchie, S. A., Lohff, A., Maldonado, C., Manley, E. A., McLin, S., Mooney, C., Ndama, M., Ogundeyi, O., Okeoma, N., Ordish, C., Padilla, N., Patrick, C., Paterson, T., Phillips, E. E., Phillips, E. M., Rampally, N., Ribeiro, C., Robert-son, M. K., Rymer, J. T., Ryan, S. M., Sammons, M., Scott, A. K., Scott, A. L., Shinomiya, A., Smith, C., Smith, K., Smith, N. L., Sobeski, M. A., Suleiman, A., Swift, J., Takemura, S., Talebi, I., Tarnogorska, D., Tenshaw, E., Tokhi, T., Walsh, J. J., Yang, T., Horne, J. A., Li, F., Parekh, R., Rivlin, P. K., Jayaraman, V., Ito, K., Saalfeld, S., George, R., Meinertzhagen, I., Rubin, G. M., Hess, H. F., Scheffer, L. K., Jain, V., and Plaza, S. M. (2020). A connectome of the adult drosophila central brain. bioRxiv.

Zhang, K. (1996). Representation of spatial orientation by the intrinsic dynamics of the head-direction cell ensemble: a theory. Journal of Neuroscience, 16(6):2112–2126.

Zhong, W., Lu, Z., Schwab, D. J., and Murugan, A. (2020). Nonequilibrium statistical mechanics of continuous attractors. Neural Computation, 32(6):1033–1068.

